# Epithelial-mesenchymal transition is the main driver of intrinsic metabolism in cancer cell lines

**DOI:** 10.1101/2021.11.02.466992

**Authors:** Sarah Cherkaoui, Stephan Durot, Jenna Bradley, Susan Critchlow, Sebastien Dubuis, Mauro Miguel Masiero, Rebekka Wegmann, Berend Snijder, Alaa Othman, Claus Bendtsen, Nicola Zamboni

## Abstract

A fundamental feature of cancer cells is genomic heterogeneity. It is a main driver of phenotypic differences, including the response to drugs, and therefore a key factor in therapy selection. Motivated by the increasing role attributed to metabolic reprogramming in tumor development, we wondered how genomic heterogeneity affects metabolic phenotype. To this end, we profiled the intracellular metabolome of 180 cancer cell lines grown in similar conditions to exclude environmental factors. For each cell line, we estimate activity for 49 pathways across the whole metabolic network. Upon clustering of activity data, we found a convergence into only two major metabolic types. These were further characterized by ^13^C-flux analysis, lipidomics, and analysis of sensitivity to perturbations. These experiments revealed differences in lipid, mitochondrial, and carbohydrate metabolism between the two major types. Finally, a thorough integration of our metabolic data with multiple omics data revealed a strong association with markers of epithelial-mesenchymal transition (EMT). Our analysis indicates that in absence of variations imposed by the microenvironment, the metabolism of cancer cell lines falls into only two major classes despite genetic heterogeneity.

## Introduction

Over the past two decades, altered metabolism has re-emerged as a prominent hallmark of cancer^1,2^. Beyond the seminal example of aerobic glycolysis^3^, multiple examples of dysregulated pathways and novel essential reactions have been presented^4–6^ and gave rise to tailored therapeutic opportunities. A key lesson in oncology that also extends to metabolism is that tumors are heterogeneous and, therefore, their sensitivities to drug or genetic treatments can differ greatly. In the context of metabolism, a main driver of heterogeneity is the tumor microenvironment. Previous studies have demonstrated the relevance of oxygenation and cancer specific nutrient utilization^7,8^ which give cancer cells a unique growth advantage. The second, intrinsic driver of heterogeneity is the genetic makeup of tumor cells, which varies between and within tumors. Mutations in coding sequences or regulatory regions and alterations in copy number may affect gene expression and the activity of proteins and, hence, enzymes. Mutations result in granular differences in pathway utilization, some of which provide a fitness advantage for tumor growth.

Even though intrinsic factors are rooted in genetic variations, genomics is poorly suited to investigate the metabolic heterogeneity of tumor cells. Beyond specific alterations that are frequently recurring in some cancer types (e.g. IDH1^6^ or PKM2^9^), sequencing of DNA or RNA fails to provide an integrated understanding of pathway activity and carbon fluxes. The latter are hard to predict because they are an emerging property governed also by nutrient availability (i.e. the microenvironment) and allosteric regulation, both of which are not captured by genomics and transcriptomics. Among the arsenal of omics technologies that are available to investigate the molecular underpinnings of cancer cells (reviewed in ^10^), metabolomics is the ideal approach to assess metabolism in action. In part, this is because regardless of the cause, intrinsic or extrinsic, changes in fluxes are associated to changes in the level of intermediates of the affected pathways. The power of metabolomics in unraveling metabolic peculiarities of cancer cells is neatly demonstrated by previous milestone studies. For instance, Jain et al.^4^ highlighted heterogeneity in metabolite uptake and secretion rates and revealed the role of glycine in tumor proliferation. Another example by Chen et al. ^11^ combined ^13^C tracing and metabolomics to reveal the relation between central carbon metabolism reprogramming and oncogenic drivers in lung cancers. More recently, Li et al.^12^ investigated the relation between intracellular metabolites and genetic alterations. They discovered the association between asparaginase’s hypermethylation and asparagine, which serves as a potential therapy for specific tumors. Overall, metabolomics has mainly been applied to the study of central carbon metabolism.

Here we leveraged the broad scope of untargeted metabolomics to characterize the intrinsic metabolic heterogeneity of a panel of 180 cancer cell lines across the full metabolic network. To exclude environmental factors, we grew the cancer cell lines in the same medium. We profiled ca. 1800 putative deprotonated metabolites across the full panel and adopted factorization by principal components to estimate activity scores for 49 metabolic pathways. In spite of the genetic variety of the tested 180 cell lines, we show that they cluster into two metabolic types, which we validated using lipidomics, ^13^C tracing, and pathway sensitivity analysis. Integration of the metabolic types with genomic and phenotypic data revealed a strong connection with EMT status, which emerges as a main determinant of intrinsic metabolic activity.

## Results

### Large-scale metabolic profiling and analysis of cancer cell lines

We set out to broadly characterize the metabolic diversity of cancer cell lines by untargeted metabolomics (Figure 1A). To gain a representative dataset, we selected a large panel of 180 cancer cell lines encompassing 11 different lineages (Figure 1B). Our panel overlapped substantially with other major cell line panels, such as the CCLE^13^, COSMIC^14^, and the NCI60^15^ (Figure 1C). As our objective was to focus on intrinsic heterogeneity and not be affected by differences in environment^16^, we cultured the cell lines in the utmost comparable condition. We grew cells in the same nutrient condition and extracted at comparable confluency and during exponential growth. Sample generation took about nine months and were organized in seven major batches. In order to assess and allow correction for batch effects, two cell lines (MCF7 and MDBADM231) were included in all batches. The study design included three biological replicates for each cell line. Each extract was analyzed twice by untargeted metabolomics by flow injection, high-resolution mass spectrometry^17^ (see details in methods). Upon data processing and quality control, the resulting metabolomic dataset including 1809 ions putatively associated to deprotonated metabolites listed in the Human Metabolome Database, for a total of 1195 measurements.

**Figure 1.**
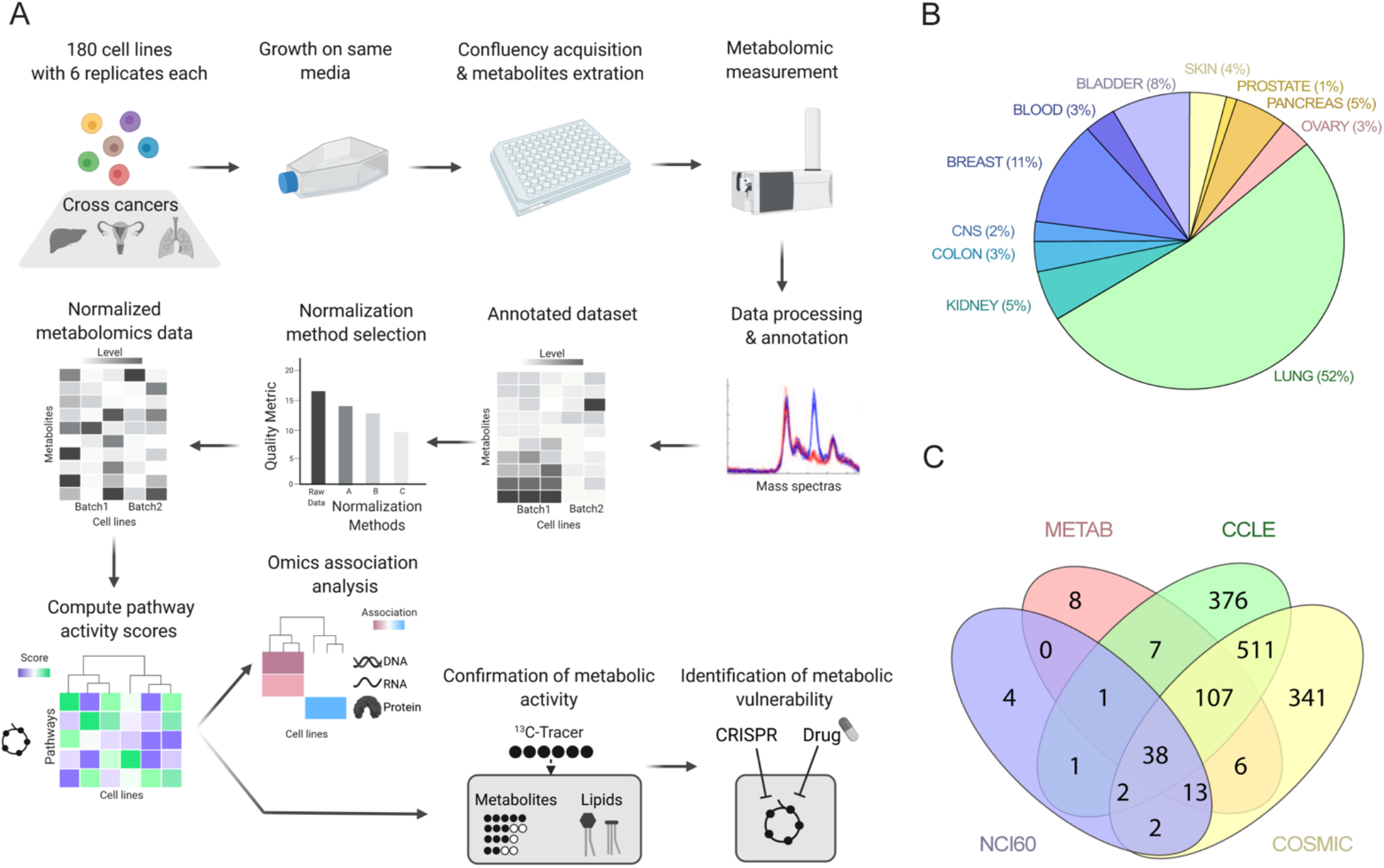
Metabolic profiling of 180 cancer cell lines. A. Schematic summarizing the workflow developed for the comparison of cancer cell lines metabolite profiles. B. 180 cancer cell lines from more than 11 tissues of origin were profiled. C. Overlap of this study cell line panel (METAB) with major cell lines resources such as Cancer Cell Line Encyclopaedia (CCLE), National Cancer Institute 60 panel (NCI60) and the Catalogue Of Somatic Mutations In Cancer panel (COSMIC).

The resulting data matrix was subject to an in-depth reproducibility analysis based on the repeated injections of MCF7 and MDBADM231 across the dataset. We used multiple quality metrics calculated for these control cell lines to measure the effect of multiple, state-of-the-art normalization procedures (see details in methods). We identified the combination of quantile normalization and ComBat^18^ as the best option to compensate for sample-to-sample differences and batch biases, respectively (Supp. Table 1).

### Inference of metabolic phenotypes

A grand challenge in analyzing metabolomics data is interpreting the cause of metabolite changes. For instance, the increase of an individual metabolite could point both to an increase of pathway flux as well as a block of the pathway immediately downstream. To distinguish between these opposite cases, it is necessary to analyze all detectable intermediates of a pathway together. In practice, when the flux of a pathway changes, a shift with coherent sign is observed for most intermediates in the pathway. This is a consequence of enzyme kinetics and the fact at that enzymes normally operate close to their substrate affinity (e.g. the Michaelis-Menten affinity constant K_M_) and far from saturation^19,20^. Therefore, flux changes cause mild but ubiquitous effects in metabolites. To capture such flux-relevant effects, we devised a strategy that uses principal component analysis to identify common trends across all detectable intermediates (see details in method). A similar concept was successfully applied to gene expression data^21^, and we demonstrate that it holds for metabolic systems with exemplary data sets^22^ (Supp. Figure 1). We first curated KEGG metabolic pathways to remove overlapping reactions across pathways. Then, for each pathway with sufficient metabolites, we projected each cell line on the first principal component, which provides a qualitative proxy for pathway activity termed ‘pathway score’. Out of the 1809 putatively annotated ions, 367 could be mapped to KEGG metabolic pathways. Based on this subset, we could infer the pathway score for 49 metabolic pathways.

To gain a top-down view of pathways scores across cancer cell lines, we applied hierarchical clustering (Figure 2). Unexpectedly, only two major clusters of cell lines emerged. The first cluster (left) was characterized with generally high activity scores for most of the pathways of central carbon metabolism, e.g. carbohydrate metabolism, amino acid metabolism, and nucleic acid metabolism. The second cluster (right) was associated with high activity scores in fewer pathways, which includes part of lipid metabolism pathways and cofactor and vitamin metabolism. In terms of pathway clustering, proximal pathways tended to group together. Pathways related to the same class also clustered together, e.g. pyrimidines, purines, or amino acids biosynthesis. Some notable exceptions remained such as tyrosine metabolism which seems to be regulated differently. Some complementary anabolic and catabolic pathway pairs had opposite activity scores. An example is fatty acid biosynthesis and degradation, which indicate that beta-oxidation and de novo lipogenesis alternate.

**Figure 2.**
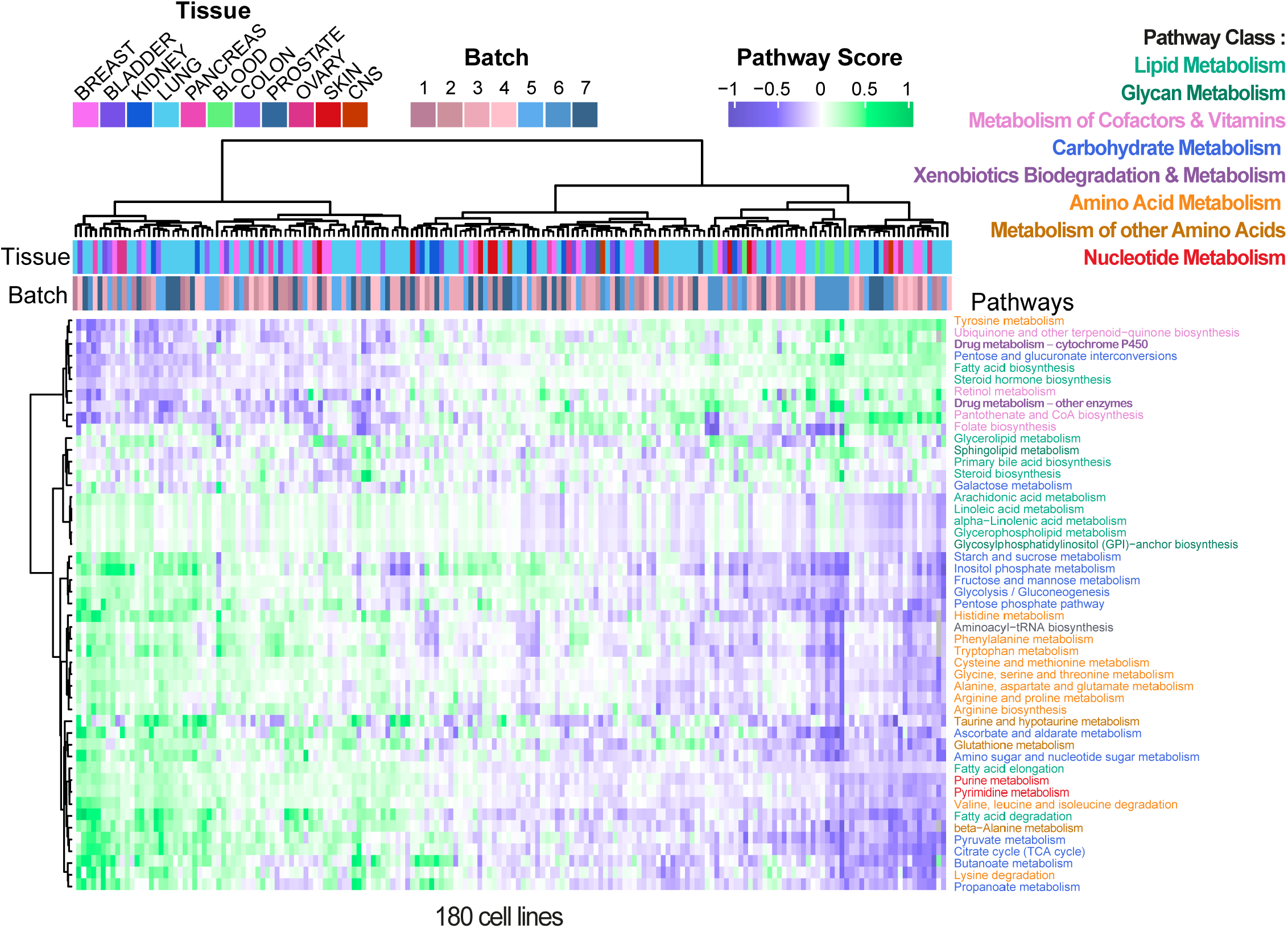
The metabolic activity of cancer cell lines. Inference of pathway score, a proxy for pathway activity, of 49 covered KEGG metabolic pathway. Hierarchical clustering of the 180 cell lines’ pathway score with pathway names colored by pathway class.

### Association of multi-omics to metabolic types

Identifying only two major clusters across a panel of 180 cancer cell lines was a surprise, because it indicates convergence into a few, robust metabolic types. Following the clustering results obtained by pathways scores, we performed an in-depth association analysis to find whether any of the tree branches was enriched in any property or trait of cell lines (e.g. tissue and batch, Figure 2). Traits included metadata related to sample collection, histologic properties, genomics, gene expression, etc. (Supp. Table 2). We assembled the information for 60’328 traits and for each calculated enrichment for all clustering subtrees with a least 18 cell lines. This resulted in 1’025’576 enrichment p-values, that were aggregated to calculate q-values that account for false discovery rate (see methods for details). By this procedure, we sought to identify the most significant associations in the clustering tree. In total, we identified 856 significant associations between a trait and a branch of the clustering derived from activity scores (Figure 3 and Supp. Table 2).

**Figure 3.**
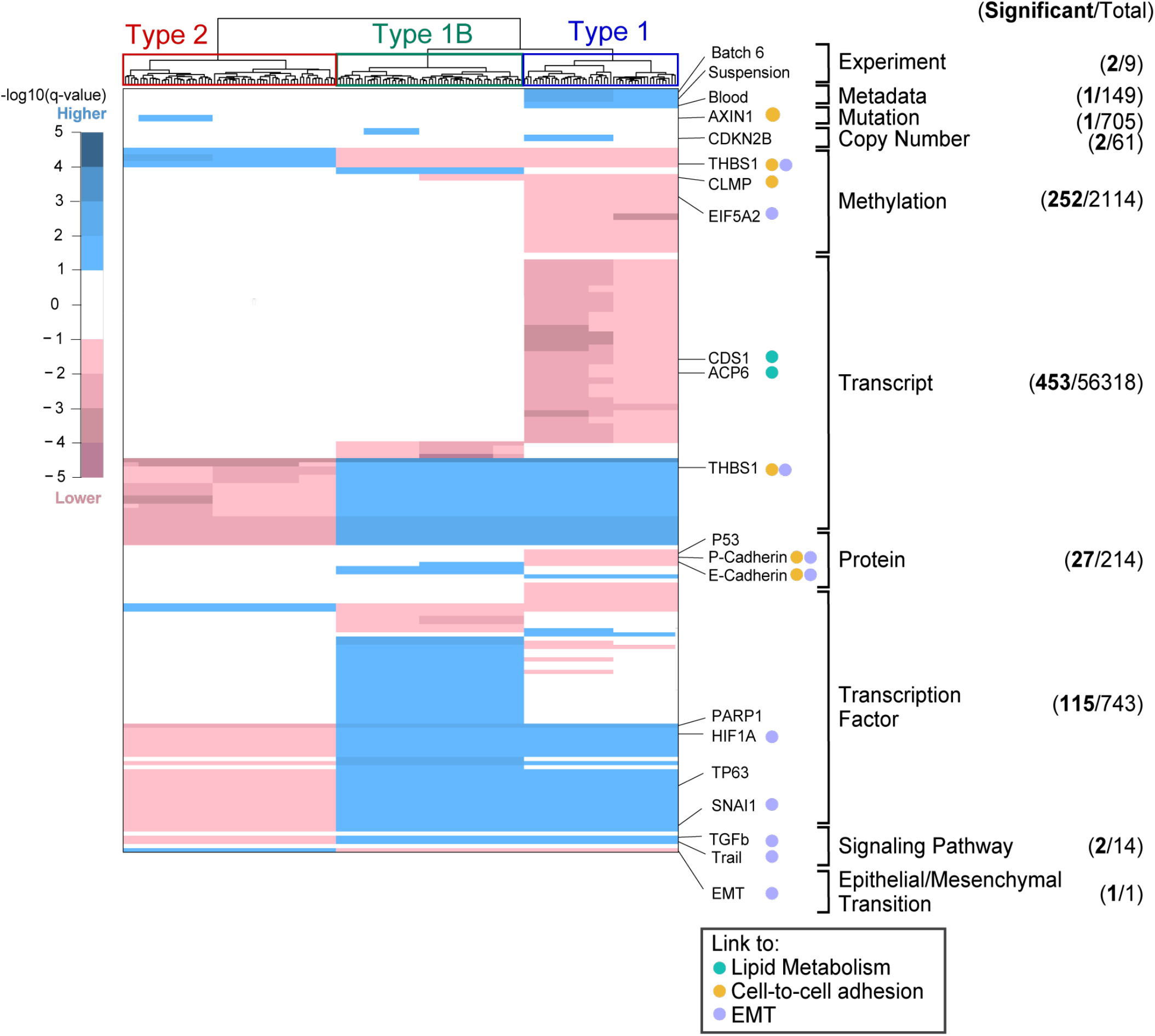
The drivers of metabolic heterogeneity. Summary of significant associations between omics and the metabolic types, displayed above as the hierarchical cluster. The list and number of traits integrated to the metabolic phenotype are listed on the right panel, where traits were considered significant at 10% FDR (cell lines n=180). For visualization, q-values were extended with a sign to indicate whether the trait is significantly higher or lower in the branch compared to the rest of the tree. Only traits significantly associated to the tree main types (1, 2, and 1B) are shown for methylation, transcript, protein and transcription. Full results are reported in data availability.

To exclude that the clustering was biased by non-cellular factors, we initially tested whether any association existed between the tree structure and experimental variables. We found that type 1, the right cluster, was associated with batch no. 6 (Figure 3). It must be noted that batch no. 6 included all cell lines that were grown in suspension. All other batches dealt with adherent cells. Rather than pointing to batch effects, we believe that this association highlights fundamental differences between growth conditions. No further association was found for any of the batches and any of the subtrees. This is a positive outcome as it indicates the clustering was not biased by experimental parameters like confluency. The enrichment analysis was extended to metadata available for many cell lines in the CCLE^23^ on cancer type, histology, histology subtype, pathology, or ethnicity of the cancer donor. In line with Li et al.^12^ and the aforementioned clustering of suspended cells, hematopoietic cells were found to be associated in the type 1. However, no further association with clinical or histological data was found.

We moved on to evaluate whether the observed metabolic types were associated with molecular traits at genomic, transcriptomic, or proteomic level. Our goal was dual: characterize major differences between metabolic types and seek potential upstream regulators that drive division in robust types. Using mutation data and focusing mainly on cancer genes^12^, we found only one association between a subcluster of type 2, the left cluster, and mutations in *AXIN1*. *AXIN1* is a component of the beta-catenin destruction complex and thus its mutation can promote the accumulation of this cell–to–cell adhesion molecule^24^. In copy number data^12^, we found two positive associations with minor subclusters. For instance, a sub-cluster of type 1 was associated with Cyclin-dependent kinase inhibitor 2B (*CDKN2b*), which acts as cell growth regulator^25^. We next considered the methylation status of cancer genes and found 252 significant associations, thereby highlighting a strong link between epigenetic regulation and metabolic phenotypes. Notable associations are highlighted, like the two negative ones between type 1 and Thrombospondin 1 (*THBS1*), and CXADR-like membrane protein (*CLMP*), both linked with cell-to-cell adhesion and interaction. *THBS1*, an activator of transforming growth factor-beta (TGF-beta), has been shown to promote an aggressive phenotype through EMT^26^. In supporting of this claim, we also found a negative association between type 1 and Eukaryotic translation initiation factor 5A-2 (*EIF5A2*), known to also induce EMT^27,28^.

In the expression data provided by the CCLE, we identified 453 associations. Because of the large number, we focused on genes that could be related to each other (identified using the String database ^29^ to find known interactions or shared biological processes among protein coding genes). Two negative associations were identified between type 1 and *CDP-diacylglycerol synthase 1 (CDS1*) and *lysophosphatidic acid phosphatase type 6* (*ACP6*). Since both target genes which encode for enzymes involved in the biosynthesis of glycerophospholipids, it may suggest a reduction in lipid synthesis in type 1. We identified positive associations between type 1 and *THBS1* expression. As previously observed, *THBS1* was less methylated in type 1 and is more expressed in type 1, hence consolidating its association to type 1. Finally, we found 27 proteins significantly associated to some subtrees. Of note, p53 levels were lower in type 1 compared to the other types. P and E-cadherin, classical markers of epithelial cells^30^ had significantly lower levels in type 1.

We hypothesized that the observed, robust metabolic types might be driven by common regulatory mechanisms. Therefore, we tested whether transcriptional factor activity and signaling pathways are associated with pathway score clusters. For 743 transcription factors (TFs), we assessed whether differential genes were overrepresented in known TF-targets^31^. We identified 115 associations between TFs and the clustered metabolic phenotypes. Several interesting hits were linked to the major types 1 and 1B, middle cluster which bears many similarity with type 1: HIF1A, previously associated with aggressive tumor phenotypes, treatment resistance, and poor clinical prognosis^32^; TP63, known to regulate migration, invasion, and *in vivo* pancreatic tumor growth^33^; and SNAI1, involved in EMT induction. Further, we tested the activity of 14 signaling pathways^34^. We identified uniquely a positive association between TGF-beta signaling and TRAIL with type 1 and 1B. Recent findings highlighted the potentially aberrant consequence of TRAIL activation in promoting cell motility and metastasis^35^ and of TGF-beta in promoting cell migration and tissue invasion^36,37^.

### Association between EMT and metabolic types

Several of the significant associations pointed to an increase of metastasis-related processes, i.e. EMT. To directly test this hypothesis, we used the EMT score proposed by Rajapakse et al.^38^. It is based on gene expression of known EMT markers to quantify the potential of invasiveness and metastasis formation of cancer. A high EMT score is associated with epithelial state and a low EMT score to mesenchymal state. In our dataset, we could confirm that type 1 and 1B were linked with the mesenchymal state, and type 2 with the epithelial state (last line, Figure 3). We validated the putative EMT association experimentally. We selected representative cell lines of the two main metabolic types 1 and 2, and stained the canonical EMT markers vimentin and E-cadherin using immunofluorescence (Supp. Figure 2). In line with the expectations, the mesenchymal marker vimentin was higher in type 1 (p-value < 1 × 10^−3^, Student t-test), and the epithelial marker E-cadherin was higher in type 2 (p-value < 1 × 10^−3^, Student t-test). Microscopy analysis also highlighted the expected morphology differences. Type 1 cells featured spindle-like shapes resembling fibroblasts, whereas type 2 portraited rounded regular shapes, consistent with EMT progression. Altogether, gene expression data, immunostaining and morphology substantiate the link between the main observed metabolic types and EMT state.

### Differences in metabolic pathway activity unraveled by ^13^C tracing

The association analysis provided novel leads on the regulatory differences that characterize the main metabolic types identified by activity scores but failed to expand our understanding of the metabolic differences. For instance, only sporadic associations were found for enzyme levels or their expression and, therefore, it is not possible to draw robust hypotheses on nutrient utilization or nutrient fluxes. To directly assess differences in pathway usage between the major metabolic types, we used ^13^C-labeling experiments. The goal was to assess whether conserved differences in fluxes could be identified between type 1 and 2. Given the generic preference of cancer cell lines for glucose and glutamine, we grew nine representative but diverse cell lines of the two types in media enriched with either [U-^13^C]glucose or [U-^13^C]glutamine for 48 hours. Upon metabolite extraction from cells, we used mass spectrometry to measure ^13^C-enrichment in metabolites and, in turn, to quantify their fraction labeling (FL), in short, their differences in ^13^C labelling. Given the experimental design in which a single substrate is labeled, the FL of each detectable metabolite informs on the fraction of carbon that originated from either glucose or glutamine. To highlight differences in carbon fluxes between type 1 and 2, we computed the difference in FL between the averages of the two groups (Figure 4A for the example of [U-^13^C]glucose). This allowed ranking all metabolites according to FL differences. To consolidate the results at the level of biochemical pathways, we sought for enrichment in both tails of the ranked metabolite list. For the example of TCA cycle metabolites, the 11 detected metabolites were mostly ranked towards type 2, resulting in a significant enrichment (q-value < 0.01, Hypergeometric test).

**Figure 4.**
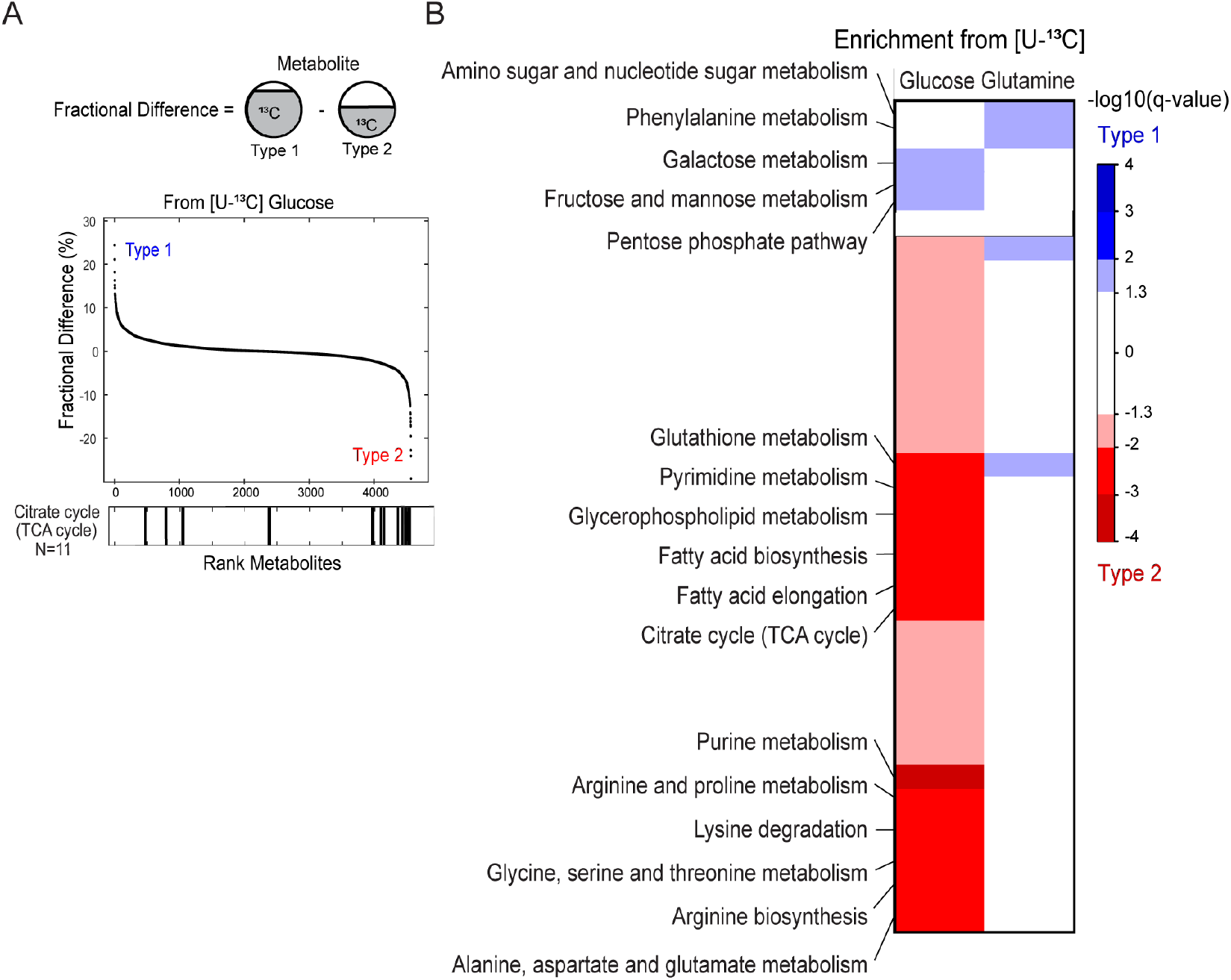
Differential pathway utilization inferred by high throughput labeling analysis. A. Schematic representation of fractional difference, difference between each metabolite fractional contribution (percentage of ^13^C labelled) between type 1 and type 2. Ranked with positive metabolite associated to type 1 and negative to type 2. Bar plot where vertical lines indicate position of metabolites of TCA cycle the fractional difference ranking. B. Metabolic pathways enriched in type 1 (blue) and type 2 (red) from [U-^13^C]glucose labelling experiment or [U-^13^C]glutamine labelling experiment (cell lines n=9, hypergeometric test corrected for FDR).

On [U-^13^C]glucose, the vast majority of pathways of primary metabolism exhibited higher ^13^C-labeling in type 2 cells (Figure 4B). This indicates that more glucose is used to replenish central carbon metabolism, amino acids, nucleotides, and fatty acids. In contrast, type 1 cells showed a slight enrichment in glucose-derived ^13^C in the pathways related to carbohydrate metabolism and storage, which are often confused because of the numerous isomers that cannot be resolved analytically. The [U-^13^C]glutamine revealed less differences between the two types, mostly because the measured FL were low in general. This indicates that glutamine-derived carbon is only a minor fraction of the total carbon assimilated for biosynthesis.

### Alterations in lipid metabolism between metabolic types

Multiple evidence suggested that the main metabolic types might differ in lipid metabolism. To validate this finding, we analyzed the lipidome of seven representative cell lines for both main types 1 and 2 by LC-MS/MS. We could detect and quantify 305 lipid species (Figure 5) and found that most lipid classes were slightly but reproducibly more abundant in type 1 cell lines (Figure 5A and B, Supp. Figure 3). Despite their minor contribution to total lipids, we stress that total cardiolipins (p-value < 0.01) and phosphatidylglycerols (p-value < 0.05) contents were higher in type 2 (Figure 5C). These lipids constitute the membrane of mitochondria and, therefore, it suggests an increase of mitochondrial mass in type 2. Conversely, lipids associated with the extracellular membrane such as phosphatidylserines (p-value < 1 × 10^−3^) and sphingomyelins (p-value < 0.01) were higher in type 1, which is consistent with the spindle-like morphology that requires increased membrane surface. The classes of ether phosphatidylcholines and triacylglycerols were characterized by remarkable within-class shifts (Supp. Figure 3). We hypothesized that these rearrangements could affect structural properties of membranes, more than the difference in total abundance. When comparing the species by double bond and acyl chain length, we found that triacylglycerols had shorter chain length in type 1 cell lines (Supp. Figure 4A) and the opposite for ether phosphatidylcholines (Supp. Figure 4B). This could point at differences in synthesis compared to uptake in these two classes. Type 2 cell lines had generally higher levels of lipid unsaturation (Supp. Figure 5), including the main constituents of the cell membrane PC (p-value < 1 × 10^−4^) and ether PC (p-value < 0.01). Lower saturation decreases tight packing of acyl chains and, in turn, increases membrane fluidity^39^.

**Figure 5.**
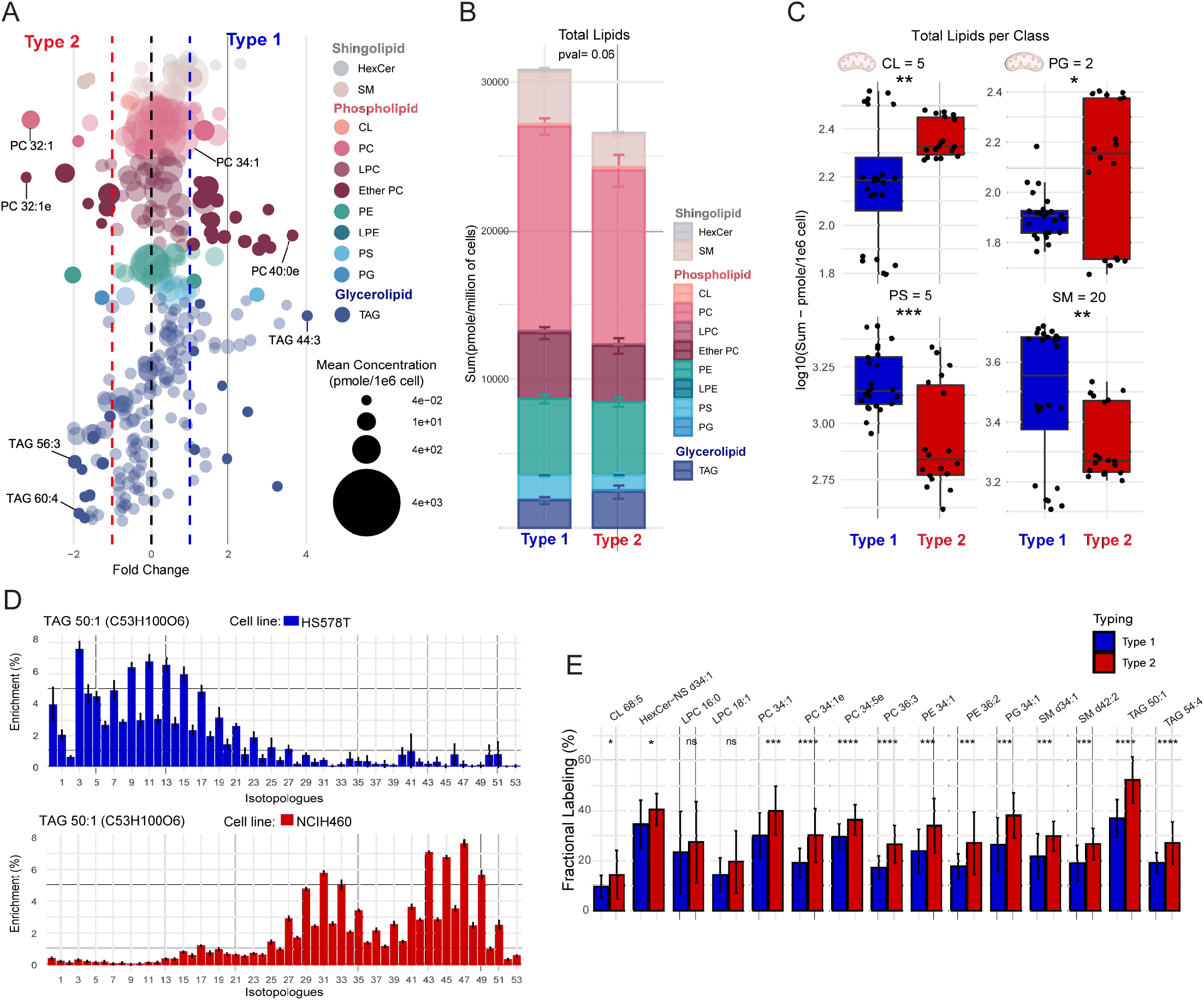
Differential lipid content and lipid biosynthesis of the metabolic types. A. Differential lipid species across type 1 and 2 in lipid class (cell lines n=7, Student t-test). B. Differences in total lipid content, with sum of average lipid class with standard error C. Differences in lipid content in each class, with lipid number in title. Lipids class linked to mitochondria identified with schema of the organelle. D. Examples of MDVs for TAG 50:1 labeled for the cell line HS578T (Type 1) and the cell line NCIH460 (Type 2) with computed average fractional labeling value (± standard deviation). E. Bar plot (mean ± standard deviation) of fractional labeling from [U-13C] glucose of most abundant lipids per lipid class (cell lines n=9, Student t-test). ns: p > 0.05, *: p <= 0.05, **: p <= 0.01, ***: p <= 0.001, ****: p <= 0.000. Abrrv: triacylglycerols (TAG), hexosyl-ceramide (HexCer), phosphatidylserine (PS), cardiolipins (CL), phosphatidylcholines (PC), phosphatidylethanolamine (PE), lysophosphatidylcholines (LPC) and sphingomyelin (SM).

Given the differences observed in lipid content between the two main metabolic types, we wondered whether we could measure differences in lipid biosynthesis rates. To assess how much of the lipids are made *de novo*, we fed cells for 48 hours with either [U-^13^C]glucose or [U-^13^C]glutamine medium. Lipid extracts were analyzed by LC-MS. Analysis of ^13^C-labeling in lipids is more challenging than for polar metabolites because of the lower abundance and the larger number of carbon atoms which causes a redistribution of the signal detected in an unlabeled sample to dozens of isotopic peaks in a labeling experiment. To maximize the quality of the data, we selected the most abundant representatives for each lipid class and performed targeted data extraction to determine full mass isotopologue distribution.

The resulting data revealed striking differences between the two major metabolic types. This is shown exemplarily for TAG 50:1 in the case of [U-^13^C]glucose, an abundant member of triacylglycerols (Figure 5D). In the type 1 representative HS578T, 24% of carbon atoms were labeled and the largest isotopologue was M+3, which results from the fusion of a ^13^C_3_-glycerol backbone and unlabeled acyl chains. In contrast, the type 2 representative NCI-H460 featured a much higher ^13^C-enrichment, 70%, with evident incorporation of ^13^C in the acyl chains. This pattern indicates substantially higher *de novo* fatty acid biosynthesis in type 2 compared to type 1. If lipogenesis is affected, similar trends should be observable across lipid classes. We extended the same analysis to nine cell lines and found for 13 out of 15 tested lipids, more ^13^C-incorporation in type 2 (p-value < 0.05, Student-test) (Figure 5E). The only exceptions being lysophosphatidylcholines, which are phospholipid derivatives and not made *de novo*. In the data related to the second tracer [U-^13^C] glutamine, the labeling enrichment was lower, in the range of 10% to 20% (Supp. Figure 6). We observed a small but opposite trend with increased ^13^C in type 1 for 5 out of tested 15 lipids which could reflect a marginal difference in the fraction of citrate that originates from glutamine and provides acetyl-CoA monomers to lipogenesis. In conclusion, we observed higher activity in *de novo* lipid synthesis (from the main carbon source glucose) in type 2 cells, which was not coupled to higher lipid content. The remaining fraction of non-labeled lipids resulted from other carbon sources, which could be from direct lipid uptake. Of note, type 2 had higher *de novo* unsaturated lipids biosynthesis and labeling into ether lipids, which are also linked with membrane fluidity^39^.

### Main metabolic types have distinct genes and drug susceptibility

To functionally validate the pathway scores, we evaluated if the inferred metabolic types were associated to differences in sensitivity to genetic or pharmacological inhibition. We used the dependency data from a CRISPR knockout screen of 18 333 genes^40^, for which 63 cell lines overlap with two types. We indeed found that active pathways were more sensitive in one type versus the other. For example, in upper glycolysis we found that PGM3 and PGAM1 deletion had stronger effect in type 1 (Student t-test p-value < 0.05 and p-value < 0.01, respectively) (Figure 6A and B). Indeed, PGAM1, *phosphoglycerate mutase 1*, knockout has a deleterious effect in both types. However, we observed that the effect was significantly more pronounced for type 1 (−1 vs −0.8 in type 2, Student t-test p-value < 0.01) and close to the median value of essential genes^40^. These two results corroborated from a functional standpoint that the association of type 1 to higher sugar metabolism activity. Inversely, the sensitivity to gene knockout shifted between major types in the TCA cycle. For example, knockout of *IDH2* or *SDHAF4* affected the growth of type 2 cells (p-value < 0.01) more than type 1.

**Figure 6.**
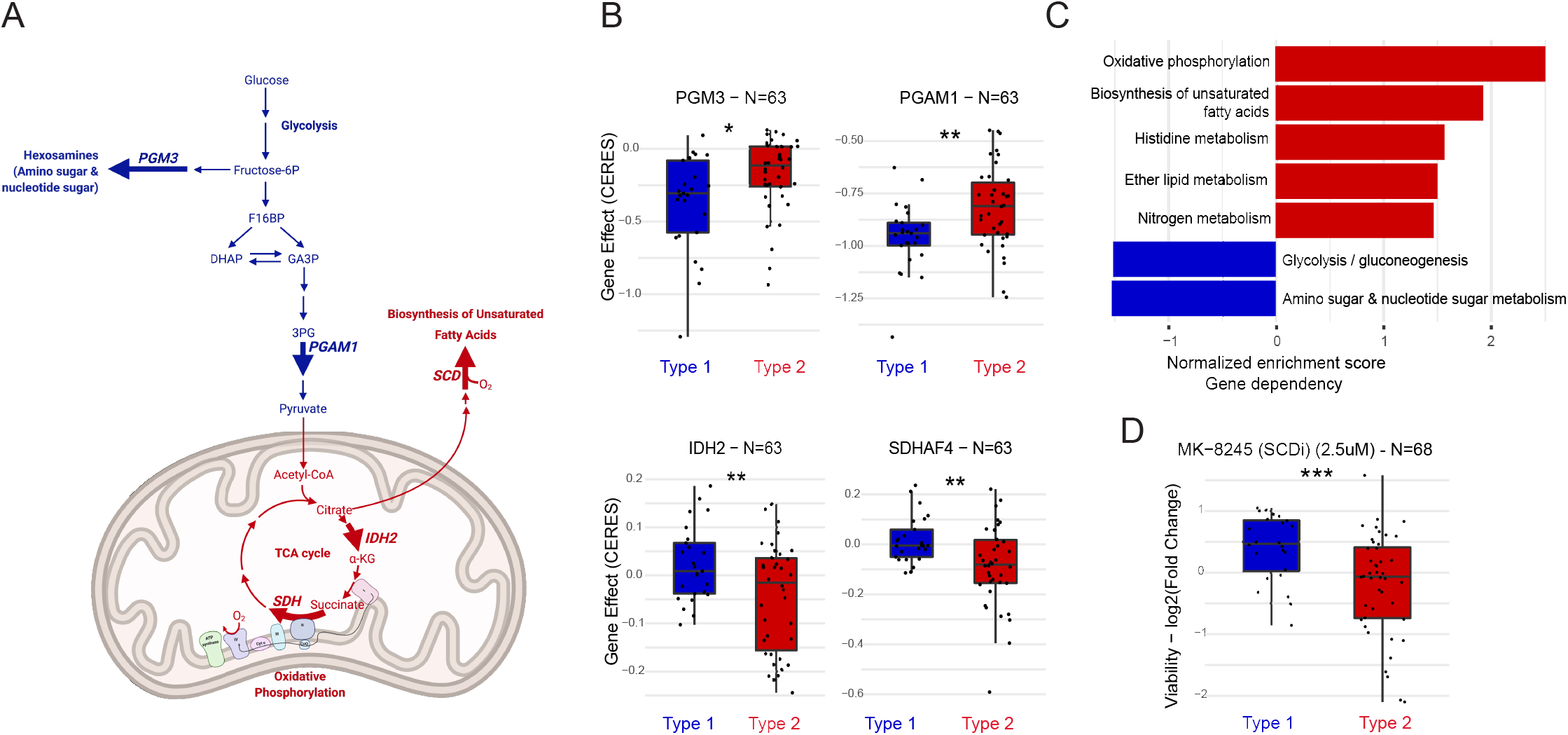
Differential sensitivity to gene dependency and drug among metabolic types. A. Schematic representation of pathways depicting reactions encoded by displayed gene dependencies. Gene dependency (CERES score) of B. PGM3, *phosphoglucomutase 3*, from the amino sugar pathway PGAM1, *phosphoglycerate mutase 1*, from glycolysis C. SDHAF4, *succinate dehydrogenase assembly factor 4 mitochondrial*, and IDH2, *isocitrate dehydrogenase 2 mitochondrial*, both from the TCA cycle (n=63, two-sided Student t-tests). C. Top pathway enrichment results computed using GSEA of the gene dependency associated to type 1 (blue) or type 2 (red). D. Drug response to MK-8245, which targets SCD, *stearoyl-CoA desaturase*, from the unsaturated fatty acids biosynthesis pathway. Gene effect (CERES score ^66^) estimate gene-dependency levels from CRISPR-Cas9 screens and is scale so that the median score of common essentials genes is −1.

We extended the analysis from single reactions to whole pathways. We ranked all 18,333 genes on their correlation with the two types and performed gene set enrichment analysis to identify pathways that include genes whose knockout lead to differential effects. The top pathways associated with type 1 were linked to sugar metabolism (Figure 6C). More strikingly, the pathways whose knock-out caused frequently a growth defect in type 2 were oxidative phosphorylation (q-value = 0, Supp. Figure 7A) and biosynthesis of unsaturated fatty acid (q-value < 0.05, Supp. Figure 7B). These are in line with the higher activity predicted by pathway score and verified by ^13^C-experiments in the TCA cycle and *de novo* lipogenesis (e.g. PC 34:5e in Figure 5E), respectively. The differentiating relevance of unsaturated fatty acids was confirmed by drug sensitivity data^41^. Type 2 cells were indeed more susceptible to inhibition of *Stearoyl-CoA desaturase* (SCD), a major contributor for biosynthesis of unsaturated fatty acids (Figure 6D). In summary, we demonstrated that the pathway activity and clustering derived from metabolomics data can translate into dependency for cancer cell lines and sensitivity to genetic or pharmacological inhibition. In type 2, the importance of mitochondrial pathways was highlighted by oxidative phosphorylation dependency and the sensitivity to unsaturated fatty acid biosynthesis was confirmed as a therapeutic liability for these cancer types.

## Discussion

We used a systematic approach to investigate the metabolic reprogramming in 180 cancer cell lines grown *in vitro* in comparable and controlled conditions. Starting from semi-quantitative data for 1809 putative deprotonated metabolites profiled by untargeted metabolomics, we estimated activity scores for 49 pathways by principal component analysis. Unsupervised clustering of cell lines based on pathway scores revealed two main groups, pointing to convergence of metabolic phenotypes. The emergence of only a few overarching groups was unexpected and, therefore, we further characterized the two major cell line metabolic types by computational and experimental means. The two metabolic types differ in pathway usage. Type 1 has enhanced carbohydrate metabolism, and type 2 relies on mitochondrial pathways, amino acid metabolism, and lipogenesis. The activity of these pathways was confirmed by isotopic labeling, and their central role was confirmed by a knockout fitness screen. Our results are coherent with the two subtypes identified by Daeman et al. in pancreatic ductal adenocarcinomas^42^ but generalized the findings to all cancer lineages and a multitude of pathways in primary and lipid metabolism.

Building on the insights from the first large-scale intracellular metabolomics study on cancer cell lines^12^, we expanded their approach of associating single genetic perturbations with single metabolites, to an approach where single perturbations are associated with global metabolic phenotypes. The integration of metabolic phenotypes to all available traits allows for the identification of possible metabolic drivers of the phenotype. Our analysis revealed many putative drivers of the major metabolic types, i.e. HIF1A, TGFbeta, and the EMT status. Their role in regulating part of metabolism is well-established^43–45^, but our work highlights how they may dominate over other regulatory axes. Albeit less common, the association between pathway activity and regulators might also reflect reversed causality, which means metabolites modulating a regulator^46^. This could be the case, for example, of the amino sugar pathway that was found to be more active in type 1. Its main product UDP-acetylglucosamine is known to affect glycosylation and, in turn, EMT^47^. Therefore, the metabolite change might act upstream of EMT, and not vice versa. Further analyses would be required to verify causality of these relations. Interestingly, the association of the types with EMT shows a much greater interaction and interconnectedness between malignant factors and metabolic reprogramming^43^.

Different from analogous studies that analyzed cancer cell lines by transcriptomics^48^, we did not find similarity in the metabolome of cell lines originating from the same lineage. Thus, the metabolic operation seems agnostic of the tumor location in the body, at least when cells are grown under similar conditions and deprived of their specific tumor microenvironment. The latter is expected to additionally impact metabolism *in vivo*, in particular if certain nutrients including oxygen become limiting^49^. This imposes additional constraints on the utilization of related pathways and increases dependence on alternative nutrients and their catabolic routes. Therefore, we expect that through action of the *in vivo* cellular environment, additional sensitivities will emerge in addition of the ones suggested by this study in rich media and normoxia^50^.

What has yet to be explored are the principles that drive these cancer cells to adopt different metabolic types. These could be associated to specific limitations that cancer cells have to bypass to support their development and transformation, such as adaptation to hypoxia, or to solve whole-cell challenges of efficient energy production or proteome allocation^51^. Because of the overwhelming result linking type 2 to aerobic pathways (TCA cycle, oxidative phosphorylation, unsaturated fatty acids, etc.), we hypothesize that oxygen and its metabolism play a major role in shaping the metabolic phenotype. Even though cells were grown under the same normoxic condition, hysteresis due to past oxygen availability could explain these phenotypes. In fact, type 1 cells are characterized by ubiquitous changes that characteristic of hypoxia: activation of HIF1 targets, inhibition of mitochondrial pathways, and increase in lipid uptake, which has been shown to be beneficial against hypoxic stress^52^. Type 2 cells, in contrast, maintain membrane fluidity by producing *de novo* unsaturated fatty acids fueled by the TCA cycle. Moreover, the differences in these aerobic pathway usages could be explained by impaired mitochondria. In fact, type 1 cells had less mitochondrial lipids, less activity in mitochondrial pathways, and were less dependent on respiration. Future work should verify causality, i.e. whether mitochondrial dysfunctions, the loss of mitochondrial mass, or stabilization of HIF1 are sufficient to drive a shift type 2 cells to type 1.

## Methods

### Cell Culture

A total of 182 cell lines were obtained and grown in seven distinct batches over the period of several months. To minimize the effect of the environment, all cell lines were adapted to growth in the same media by maintaining them in culture for at least 2 weeks. Cells were cultured in RPMI 1640 Phenol Red Free (Sigma #R7059) with 2 mM L-glutamine (Sigma #G7513, Lot #RNBD0904) freshly added. The medium was supplemented with 10% fetal calf serum (PAA Laboratories, Linz, Austria, #A15-701, Lot #A30111-3524) and is referred hereafter as RPMI, 10% FCS, L-glutamine. Cell lines were grown at 37 °C with 5% CO_2_. Each cell line was seeded into 3 wells of 6-well plates and grown for 48 hours. Cell lines were maintained according to standard protocols, their identity verified via STR sequencing and tested for mycoplasma infections. All adherent cells were grown to reach similar confluency. The cut-off values for adherent cell lines are a minimum of 50%, and a maximum of 80% confluency. For mixed suspension/adherent lines, they were judged on a case-by-case basis, and the minimum confluency as it does not reflect direct cell density set at 35%. For suspension cells, after 48h, cell culture counts were accessed in the count plate on an automated cell counter (Cedex, by Roche, Basel, Switzerland). As there was little or no growth lag after splitting the culture, we calculated growth rates from recent counts and seeded accordingly. The final density was at the upper end of the exponential phase of growth.

### Metabolite extraction

At 48h, the medium was removed via aspiration, and cells were washed twice with a wash solution (75 mM Ammonium Carbonate, adjusted to pH 7.4 with Acetic Acid). Metabolites were quenched by dipping the bottom of the plate in liquid nitrogen for 1 min. Metabolites were extracted using a 40% methanol, 40% acetonitrile, 20% water solvent. The plate was sealed and incubated at −20 °C for, 10 min. Extracted cells were scraped off the bottom of each well using a pipet with wide-bore tips. Next, the cell extracts were transferred to 96-well plates with conical bottom and centrifuged at 4 °C, 2800rpm for 30 min to separate cell debris. The cleared supernatants were injected for mass spectrometric analysis.

### Untargeted metabolomics

Untargeted metabolite profiling was performed using flow injection analysis on an Agilent 6550 QTOF instrument (Agilent, Santa Clara, CA) using negative ionization, 4 GHz high resolution acquisition, and scanning in MS1 mode between m/z 50-1000 at 1.4 Hz^17^. The solvent was 60:40 isopropanol:water supplemented with 1 mM NH_4_F at pH 9.0, as well as 10 nM hexakis(1H, 1H, 3H-tetrafluoropropoxy)phosphazine and 80 nM taurochloric acid for online mass calibration. The seven batches were analyzed sequentially. Within each batch, the injection sequence was randomized. Data was acquired in profile mode, centroided and analyzed with Matlab (The Mathworks, Natick). Missing values were filled by recursion in the raw data. Upon identification of consensus centroids across all samples, ions were putatively annotated by accurate mass and isotopic patterns. Starting from the HMDB v3.0 database^53^, we generated a list of expected ions including deprotonated, fluorinated, and all major adducts found under these conditions. All formulas matching the measured mass within a mass tolerance of 0.001 Da were enumerated. As this method does not employ chromatographic separation or in-depth MS2 characterization, it is not possible to distinguish between compounds with identical molecular formula. The confidence of annotation reflects Level 4 but – in practice - in the case of intermediates of primary metabolism it is higher because they are the most abundant metabolites in cells. The resulting data matrix included 1809 ions that could be matched to deprotonated metabolites listed in HMDB v3.0. All m/z peaks that remained unmatched or were associated to adducts or heavy isotopomers were discarded.

### Data normalization

As first-line quality control for each measurement, we computed the sum of all intensities (TIC) of each injection. If extremely low or high, the TIC is a sign of problems in the sample preparation, injections, or measurement. Out of the 1387 injections, 68 exhibited abnormal TIC and were removed. These included all six replicates of PC14PE6 and NCIH446. Out of the initial 182 cell lines included in the screen, we were left with data for 180. Large-scale, untargeted metabolomics experiments are naturally subject to various factors that introduce unwanted effects on data: batch effects of sampling and analysis, drifts in measurements, accumulation of dirt, drops in instrument sensitivity, etc. To address these potential issues, we tested numerous normalization procedures. To assess the efficacy of each procedure, we adopted quantitative measures of reproducibility. The fundament of the reproducibility analysis was the inclusion of the two cell lines as quality controls (QC): MCF7 and MDAMB321 in each of the seven batches. Because of sequential analysis of batches, representatives of both cell lines were distributed over the entire injection sequence. The expectation was that all repeated measurements of either MCF7 or MDAMB321 had to be possibly similar, but the differences between MCF7 and MDAMB321 were preserved across the seven batches. The reproducibility metrics included the following criteria:

- Batch Scoring fold change (FC): calculated as the mean fold-change between random subsets of MCF7 cell line measurements. Out of the 66 replicate measurements for this cell lines across all batches, we randomly sampled two sets of 6 samples, calculated the mean intensities for each set and, in turn, the fold-change for each of the 1809 metabolites ions. The sampling was repeated 1000 times. From the distribution of all fold-changes, we calculated the QC distance as the threshold that corresponds to the 5% false discovery rate. Ideally, the Batch Scoring FC between replicates of the same cell line should be zero. Normalization improves reproducibility if the threshold value decreases.
- FC Reproducibility: for each batch, we calculated the fold-change for all 1809 metabolite ions between the MCF7 and MDAMB321 replicates. We calculated the mean Euclidean distance between the fold-changes of all seven batches. Normalization improves reproducibility if the distance between fold-changes decreases.
- FC Reproducibility amino acids (AA): for each batch, we calculated the fold-change between the MCF7 and MDAMB321 replicates for ions feature matching to AA. We calculated the mean Euclidean distance between the fold-changes of all seven batches. Normalization improves reproducibility if the distance between fold-changes decreases.
- Interbatch Distance: we adopted the metric defined by Wehrens at al. ^54^. It slices the data by batches, applies principal component analysis to each batch, and computes the Bhattacharyya distance. Normalization improves reproducibility if the distance decreases.
- Kolmogorov-Smirnov: we tested for ions whether MCF7 samples from different batches were on the same distribution. A two-sample Kolmogorov-Smirnov was used to test for this batch effect. We counted the frequency of tests that revealed batch effects, by counting the number of tests significant (p-value < 0.05) on the total number of tests done.

The five criteria were scaled across all raw and normalized datasets to [0…1] and finally averaged to obtain a single reproducibility score.

We tested 15 different normalization procedures including state-of-the-art methods. Conceptually, these methods can be divided into 3 different classes:

i. Methods that correct for user-defined batches. In our case, we used the seven experimental batches. We tested two implementations of the Remove Unwanted Variation (RUV) method^55^ and the empirical Bayes method ComBat^56^. In some cases, the MDAMB321 samples were selected as QC samples to be used for correction.
ii. Sample variance: as a consequence of differences in cell amount, sampling, pipetting errors, injection errors, etc. variations can occur in single samples. To correct for such issues, we tested normalization using quantiles (Quantile), log10-mean (MEAN), median (MEDIAN), standard deviation (STD), median absolute deviation (MAD), the sum of all ions (TIC), probabilistic quotient (PQN), and scaling with cellular confluency at the time of sampling (Confluency).
iii. Methods that correct for signal drifts, that occur chronologically during the injection sequence. The drifts might be caused by smooth changes in solvents, the ionization source, ion optics, or the detection process. We used three methods that consider the injection sequence to detect temporal drifts and correct after smooth interpolation. First, we implemented a method that applies a moving median (window 120 min) to all measured samples to estimate a robust trendline (MovMed). Second, we used a locally weighted regression (LOESS)^57^ and its derivate for temporal trends (Robust LOESS)^58^, and third, we used the QC-based support vector regression method (QC-SVR)^59^. In the latter case, the MDAMB321 samples were selected as QC samples to be used for correction.

These methods were tested singularly and in reasonable combinations (Combo). Given that the three classes tackle different types of problems, we have combine up to methods from the different classes. The choice of methods and the order of combinations was based on their improvement of quality metric scores. Results and average quality score can be found in Supp. Table 1.

### Metabolic pathway definition

A common issue using pathways is that their definition can be arbitrary, i.e. the start and end of a pathway is dependent on the database. Kyoto Encyclopedia Genes and Genomes (KEGG)^60^, our chosen database because of its high curation, has the disadvantage of having substantially overlapping pathways. We circumvented this limitation by removing reactions, and their corresponding substrate or product, which were present in multiple metabolic pathways. This curation resulted in a smaller pathway definition with less overlapping reactions, and thus metabolites more specific to a pathway. Out of the 1809 putatively annotated ions (according to HMDB), 367 could be linked to KEGG pathways. As in many cases the measurement doesn’t not allow to distinguish between structural isomers, an ion could match to one or multiple metabolites with the same formula. In total, the 367 deprotonated ions matched to 530 metabolites which are part of KEGG metabolic pathways.

### Pathway activity scoring

Our goal is to infer flux changes from relative metabolite abundances, one pathway at the time. At biochemical level, the relation between metabolites and fluxes is governed by enzyme kinetics and capacity. If pathway flux change, we expect all intermediate concentrations to shift in a coherent direction. In some cases, the metabolite changes may only be subtle, depending on enzyme saturation and the additional regulatory mechanisms. In general, coherent and distributed changes of pathway metabolites indicate an underlying flux change. In contrast, strong but local metabolite changes reflect pathway interruption^61^ or changes in enzyme kinetics or abundance that are compensated by a modulation of the reactants but don’t influence pathway flux^62^. Hence, we sought to quantify ubiquitous and coordinated changes in multiple metabolites within each pathway. This was done by principal component analysis (PCA). PCA identifies components which best capture metabolite variance. As we are interested in the major metabolite effect, we focused on the first principal component (PC1) and used the PC1 scores as proxy of pathway activity. The same principle has been adopted in the past for the analysis of transcriptomics data^21^. To verify the validity of the method in the case of metabolomics, we used a published dataset by Hackett et al. ^22^which offered both metabolomics and measured fluxes for multiple conditions with sufficiently diverse flux distributions (Supp. Figure 1). For representative pathways, we correlated PC1 scores and measured fluxes. Strong positive correlation is observed between the glycolytic and pentose phosphate metabolites summarized in PC1 and the fluxes of these pathways. As the direction of principal components can be flipped (e.g. in the case of the purine pathway in Supp. Figure 1C). Overall, the PC1 scores correlated favorably with fluxes in all cases tested.

### Inference of pathway score in cancer cell lines

Metabolomics data were mapped onto pathways, where pathways were considered with a minimum of four metabolites measured were further analyzed. Regardless of the number of detectable metabolites, the relative pathway score for each cell line replicate was obtained by PCA as outlined above. To isolate robust pathway scores, we analyzed the 1060 injections (after removal of controls) independently. For each cell line, we averaged the 6 independent PC1 scores to assess the pathway activity score. Final scores were scaled to [−1…1] for comparison across pathways.

### Typing and association analysis

The matrix with pathway scores was subject to hierarchical clustering using Ward’s method.

The association analysis over the tree was done by iterating through all subtrees with at least 18 cell lines (10% of the total number). For categorical traits (e.g. batch number or genomic data), we calculated enrichments with hypergeometric tests. For continue variables (e.g. gene expression), we used Student two-tailed t-tests. We assembled all (1 025 576) resulting p-values and corrected *in toto* for false discovery rate by the Storey & Tibshirani method to produce q-values^63^.

### Selection of cell lines for follow ups

We chose 9 cells from type 1 and type 2 for further evaluation (Supp. Table 3). These cell lines were selected to span diverse lineages and growth rates. To show differences in metabolism across the same lineage we selected pairs of ovarian cell lines (OVCAR3 and OVCAR5) and of breast cell lines (T47D and HS578T) belonging to different types. To confirm that the metabolic types are not associated with growth rate, we selected cell lines with doubling time spanning from 17h to 53h and mixed type. Cell lines were obtained from the National Cancer Institute (NCI, Bethesda, MD, USA). After thawing, the cell lines were expanded in cell culture flasks (Nunc T75, Thermo Scientific) at 37°C and 5% CO2 in RPMI-1640 (Biological Industries, cat.no. 01-101-1A) supplemented with 5% fetal bovine serum (FBS, Sigma Aldrich, cat.no. F6178), 2 mM L-glutamine (Gibco, cat.no. 25030024), 2 g/L D-glucose (Sigma Aldrich, cat.no. G8644), and 100 U/mL penicillin/streptomycin (P/S, Gibco, cat.no. 15140122).

### Cell imaging and image analysis

Cells were stained overnight at 4°C in blocking solution with the following antibodies: AlexaFluor® 488 anti-Vimentin (1:1000, Biolegend, #677809), AlexaFluor® E-cadherin (1:200, clone 36/E-cadherin, BD #560062), and DAPI (10mg/ml, 1:1000, Sigma Aldrich). High-content imaging was performed with an Opera Phenix automated spinning-disk confocal microscope at 40x magnification (Perkin Elmer, HH14000000). To measure cell area shape features, single cells were segmented using CellProfiler 2.2.063. Nuclei segmentation relied on the DAPI channel. CellProfiler module ‘DetectPrimaryObject’ was used to identify the nuclei and ‘DetectSecondaryObject’ was used to derive the intensity of the marker in the area around the nucleus. For each cell, 9 images from 9 biological replicates were segmented, which cumulated into 10 273 segmented cells. Statistical significance was assessed by Student’s t-test.

### ^13^C labeling experiments

After two passages, FBS in the growth medium was replaced by dialyzed FBS with a reduced content of low molecular weight compounds (dFBS, Sigma Aldrich, cat.no. F0392). 3 replicates of each cell line were grown for 48h in either growth medium with either [U-^13^C] glucose (Sigma-Aldrich, cat.no. 389374), [U-^13^C] glutamine (Cambridge Isotope Laboratories, cat.no. CLM-1822-H-PK), or naturally labeled growth medium. At 48h, we extracted metabolites as described and analyzed the extracts by LC-MS on an Agilent 6546 Q-TOF instrument (Agilent, Santa Clara, CA). The liquid chromatography consisted of a 30 mm Waters ACQUITY UPLC BEH C18 column (cat. no. 186002352) and a linear gradient from 10:90 to 90:10 (v/v) water:methanol mix. Mass spectra were recorded from 50 to 1050 m/z in 4 GHz HighRes, negative ionization mode. Annotation was done by matching the measured mass of the ions with reference compounds derived from the Human Metabolome Database (HMDB 4.0), taking labeling patterns of potential metabolites into consideration.

### Lipidomics

Cells were grown in the same three conditions as above: naturally labeled, [U-^13^C] glucose, and [U-^13^C] glutamine medium. At 48h, internal standard (EquiSPLASH, Avanti Polar Lipids, cat.no. 330731) was added to all cell lines to enable the quantification of lipid species. Lipid extraction was performed using 50:50 (v/v) methanol/isopropanol for 1 h at −20°C. Untargeted lipidomics was performed by LC-MS on a Thermo Fisher Q-Exactive HF-X mass spectrometer (Thermo, Massachusetts, United States). For liquid chromatography we used a 30 mm Waters ACQUITY UPLC BEH C18 column (cat. no. 186002352) and a 7 minute gradient from 15% buffer B (90% (v/v) isopropanol, 10% acetonitrile, 10mM ammonium acetate) and 85% buffer A (60% acetonitrile, 50% water, 10mM ammonium acetate) to 99% Buffer B. Mass spectra were recorded from 150 to 2000 m/z in positive ionization mode, recording MS1 and MS2 (DDA, top 5 ions) spectra. Lipidomics data processing for non-labeled samples was done using Compound Discoverer 3.1 (Thermo, Massachusetts, United States). Lipid were annotated with MS2 information. Lipids from each class were quantified using class-specific internal standards.

For labeled lipids, we adopted a targeted data extraction. The most abundant representatives for each lipid class were selected in naturally labeled samples. All ^13^C-isotopomer traces were extracted as ion chromatograms from labeled samples based on accurate mass and retention time. Related mass isotopomers were integrated with identical boundaries and normalized to unity to obtain labeling fractions.

### Gene dependency and drug response analysis

Gene dependencies were obtained from Tsherniak et al.^40^ and the response from Corsello et al.^41^. Gene set enrichment was performed using GSEA (https://www.gsea-msigdb.org/gsea/index.jsp) leading-edge analysis^65^ by correlating gene dependency to the two types. Gene sets were taken from the curated KEGG pathways described above. We used 1000 permutations of the gene-level values to calculate normalized enrichment scores and statistical significance. GSEA results display the enrichment score normalized to mean enrichment of random samples of the same size.

## Acknowledgements

We thank Mattia Zampieri for providing the cell lines for the follow up experiments, the transcription factor dataset, and for his advice. We thank Julio Saez-Rodriguez, Aurélien Dugourd, Enio Gjerga and Panuwat Trairatphisan for providing the signaling pathway dataset and for their advice. We thank Jonathan DeGeer for his comments on the manuscript.

## Data availability

Raw metabolomics files of the 180 cancer cell lines can be accessed from the Massive database (https://massive.ucsd.edu/ProteoSAFe/dataset.jsp?accession=MSV000087155). Data tables and raw files of follow up experiments are available at https://doi.org/10.3929/ethz-b-000511784.

## References

1. Ward, P. S. & Thompson, C. B. Metabolic Reprogramming: A Cancer Hallmark Even Warburg Did Not Anticipate. Cancer Cell vol. 21 (2012).

2. Pavlova, N. N. et al. The Emerging Hallmarks of Cancer Metabolism. Cell Metabolism 23, 27–47 (2016).

3. Warburg, O. & Minami, S. Versuche an Überlebendem Carcinom-gewebe. Klinische Wochenschrift 2, (1923).

4. Jain, M. et al. Metabolite Profiling Identifies a Key Role for Glycine in Rapid Cancer Cell Proliferation. Science 336, 1040–1044 (2012).

5. Son, J. et al. Glutamine supports pancreatic cancer growth through a KRAS-regulated metabolic pathway. Nature 2013 496:7443 496, 101–105 (2013).

6. Dang, L. et al. Cancer-associated IDH1 mutations produce 2-hydroxyglutarate. Nature 462, 739–744 (2009).

7. DeBerardinis, R. J. et al. Beyond aerobic glycolysis: Transformed cells can engage in glutamine metabolism that exceeds the requirement for protein and nucleotide synthesis. Proceedings of the National Academy of Sciences of the United States of America 104, 19345–19350 (2007).

8. Spinelli, J. B. et al. Metabolic recycling of ammonia via glutamate dehydrogenase supports breast cancer biomass. Science 358, 941–946 (2017).

9. Dayton, T. L., Jacks, T. & Heiden, M. G. vander. PKM2, cancer metabolism, and the road ahead. EMBO reports 17, 1721–1730 (2016).

10. Li, S., Zhang, Z. & Han, L. Molecular Treasures of Cancer Cell Lines. Trends in Molecular Medicine 25, 657–659 (2019).

11. Chen, P.-H. et al. Metabolic Diversity in Human Non-Small Cell Lung Cancer Cells. Molecular cell 76, 838–851 (2019).

12. Li, H. et al. The landscape of cancer cell line metabolism. Nature Medicine 25, 850–860 (2019).

13. Ghandi, M. et al. Next-generation characterization of the Cancer Cell Line Encyclopedia. Nature 569, 503–508 (2019).

14. Forbes, S. A. et al. COSMIC: exploring the world’s knowledge of somatic mutations in human cancer. Nucleic acids research 43, D805–11 (2015).

15. Shoemaker, R. H. The NCI60 human tumour cell line anticancer drug screen. Nature Reviews Cancer 6, 813–823 (2006).

16. Lagziel, S., Gottlieb, E. & Shlomi, T. Mind your media. Nature Metabolism 2020 2:12 2, 1369–1372 (2020).

17. Fuhrer, T., Heer, D., Begemann, B. & Zamboni, N. High-throughput, accurate mass metabolome profiling of cellular extracts by flow injection-time-of-flight mass spectrometry. Analytical chemistry 83, 7074–80 (2011).

18. Johnson, W. E., Li, C. & Rabinovic, A. Adjusting batch effects in microarray expression data using empirical Bayes methods. Biostatistics 8, (2007).

19. Park, J. O. et al. Metabolite concentrations, fluxes and free energies imply efficient enzyme usage. Nature chemical biology 12, 482–9 (2016).

20. Bar-Even, A. et al. The Moderately Efficient Enzyme: Evolutionary and Physicochemical Trends Shaping Enzyme Parameters. Biochemistry 50, 4402–4410 (2011).

21. Segura-Lepe, M. P., Keun, H. C. & Ebbels, T. M. D. Predictive modelling using pathway scores: robustness and significance of pathway collections. BMC Bioinformatics 20, 543 (2019).

22. Hackett, S. R. et al. Systems-level analysis of mechanisms regulating yeast metabolic flux. Science 354, (2016).

23. Ghandi, M. et al. Next-generation characterization of the Cancer Cell Line Encyclopedia. Nature 569, 503–508 (2019).

24. Jeong, W.-J., Ro, E. J. & Choi, K.-Y. Interaction between Wnt/β-catenin and RAS-ERK pathways and an anti-cancer strategy via degradations of β-catenin and RAS by targeting the Wnt/β-catenin pathway. npj Precision Oncology 2, (2018).

25. Adamovic, T. et al. Rearrangement and allelic imbalance on chromosome 5 leads to homozygous deletions in the CDKN2A/2B tumor suppressor gene region in rat endometrial cancer. Cancer Genetics and Cytogenetics 184, (2008).

26. Jayachandran, A. et al. Thrombospondin 1 promotes an aggressive phenotype through epithelial-to-mesenchymal transition in human melanoma. Oncotarget 5, (2014).

27. Zhu, W. et al. Overexpression of EIF5A2 promotes colorectal carcinoma cell aggressiveness by upregulating MTA1 through C-myc to induce epithelial - mesenchymaltransition. Gut 61, (2012).

28. Khosravi, S. et al. Role of EIF5A2, a downstream target of Akt, in promoting melanoma cell invasion. British Journal of Cancer 110, (2014).

29. Szklarczyk, D. et al. The STRING database in 2017: Quality-controlled protein-protein association networks, made broadly accessible. Nucleic Acids Research 45, (2017).

30. Ribeiro, A. S. & Paredes, J. P-cadherin linking breast cancer stem cells and invasion: A promising marker to identify an “intermediate/metastable” EMT state. Frontiers in Oncology 4, (2015).

31. Ortmayr, K., Dubuis, S. & Zampieri, M. Metabolic profiling of cancer cells reveals genome-wide crosstalk between transcriptional regulators and metabolism. Nature Communications 10, (2019).

32. Wigerup, C., Påhlman, S. & Bexell, D. Therapeutic targeting of hypoxia and hypoxia-inducible factors in cancer. Pharmacology and Therapeutics vol. 164 (2016).

33. Somerville, T. D. D. et al. TP63-Mediated Enhancer Reprogramming Drives the Squamous Subtype of Pancreatic Ductal Adenocarcinoma. Cell Reports 25, (2018).

34. Schubert, M. et al. Perturbation-response genes reveal signaling footprints in cancer gene expression. Nature Communications 9, (2018).

35. Fulda, S. The dark side of TRAIL signaling. Cell Death and Differentiation vol. 20 (2013).

36. Yeh, H. W., Lee, S. S., Chang, C. Y., Lang, Y. D. & Jou, Y. S. A new switch for TGFβ in cancer. Cancer Research 79, (2019).

37. Hao, Y., Baker, D. & ten Dijke, P. TGF-β-Mediated Epithelial-Mesenchymal Transition and Cancer Metastasis. International journal of molecular sciences 20, (2019).

38. Rajapakse, V. N. et al. CellMinerCDB for Integrative Cross-Database Genomics and Pharmacogenomics Analyses of Cancer Cell Lines. iScience 10, 247–264 (2018).

39. Jain, I. H. et al. Genetic Screen for Cell Fitness in High or Low Oxygen Highlights Mitochondrial and Lipid Metabolism. Cell 181, 716–727.e11 (2020).

40. Tsherniak, A. et al. Defining a Cancer Dependency Map. Cell 170, 564–576.e16 (2017).

41. SM, C. et al. Discovering the anti-cancer potential of non-oncology drugs by systematic viability profiling. Nature Cancer 1, 235–248 (2020).

42. Daemen, A. et al. Metabolite profiling stratifies pancreatic ductal adenocarcinomas into subtypes with distinct sensitivities to metabolic inhibitors. Proceedings of the National Academy of Sciences 112, E4410–E4417 (2015).

43. Georgakopoulos-Soares, I., Chartoumpekis, D. v., Kyriazopoulou, V. & Zaravinos, A. EMT Factors and Metabolic Pathways in Cancer. Frontiers in Oncology vol. 10 499 (2020).

44. Hua, W., ten Dijke, P., Kostidis, S., Giera, M. & Hornsveld, M. TGFβ-induced metabolic reprogramming during epithelial-to-mesenchymal transition in cancer. Cellular and Molecular Life Sciences 2019 77:11 77, 2103–2123 (2019).

45. Semenza, G. L. Hypoxia-Inducible Factors in Physiology and Medicine. Cell 148, 399–408 (2012).

46. Jia, D. et al. Towards decoding the coupled decision-making of metabolism and epithelial-to-mesenchymal transition in cancer. British Journal of Cancer 2021 124:12 124, 1902–1911 (2021).

47. Akella, N. M., Ciraku, L. & Reginato, M. J. Fueling the fire: emerging role of the hexosamine biosynthetic pathway in cancer. BMC Biology 17, 52 (2019).

48. Hu, J. et al. Heterogeneity of tumor-induced gene expression changes in the human metabolic network. Nature Biotechnology 31, 522–529 (2013).

49. Hensley, C. T. et al. Metabolic Heterogeneity in Human Lung Tumors. Cell 164, 681–694 (2016).

50. Eylem, C. C. et al. Untargeted multi-omic analysis of colorectal cancer-specific exosomes reveals joint pathways of colorectal cancer in both clinical samples and cell culture. Cancer Letters 469, 186–194 (2020).

51. Basan, M. et al. Overflow metabolism in Escherichia coli results from efficient proteome allocation. Nature 528, (2015).

52. Kamphorst, J. J. et al. Hypoxic and Ras-transformed cells support growth by scavenging unsaturated fatty acids from lysophospholipids. Proceedings of the National Academy of Sciences of the United States of America 110, (2013).

53. Wishart, D. S. et al. HMDB 3.0--The Human Metabolome Database in 2013. Nucleic acids research 41, D801–7 (2013).

54. Wehrens, R. et al. Improved batch correction in untargeted MS-based metabolomics. Metabolomics 12, 88 (2016).

55. Risso, D., Ngai, J., Speed, T. P. & Dudoit, S. Normalization of RNA-seq data using factor analysis of control genes or samples. Nature Biotechnology 32, 896–902 (2014).

56. Johnson, W. E., Li, C. & Rabinovic, A. Adjusting batch effects in microarray expression data using empirical Bayes methods. Biostatistics 8, (2007).

57. Cleveland, W. S. & Devlin, S. J. Locally weighted regression: An approach to regression analysis by local fitting. Journal of the American Statistical Association 83, 596–610 (1988).

58. Cleveland, W. S. Robust locally weighted regression and smoothing scatterplots. Journal of the American Statistical Association 74, 829–836 (1979).

59. Kuligowski, J., Sánchez-Illana, Á., Sanjuán-Herráez, D., Vento, M. & Quintás, G. Intra-batch effect correction in liquid chromatography-mass spectrometry using quality control samples and support vector regression (QC-SVRC). Analyst 140, 7810–7817 (2015).

60. Kanehisa, M. & Goto, S. KEGG: kyoto encyclopedia of genes and genomes. Nucleic acids research 28, 27–30 (2000).

61. Fuhrer, T., Zampieri, M., Sévin, D. C., Sauer, U. & Zamboni, N. Genomewide landscape of gene–metabolome associations in Escherichia coli. Molecular Systems Biology 13, 907 (2017).

62. Fendt, S. M. et al. Tradeoff between enzyme and metabolite efficiency maintains metabolic homeostasis upon perturbations in enzyme capacity. Molecular Systems Biology 6, (2010).

63. Storey, J. D. & Tibshirani, R. Statistical significance for genomewide studies. Proceedings of the National Academy of Sciences 100, 9440–9445 (2003).

64. McQuin, C. et al. CellProfiler 3.0: Next-generation image processing for biology. PLOS Biology 16, e2005970 (2018).

65. Subramanian, A., Tamayo, P., Mootha, V. K., Mukherjee, S. & Ebert, B. L. Gene set enrichment analysis : A knowledge-based approach for interpreting genome-wide. Proceedings of the National Academy of Sciences 43, 15545–50 (2005).

66. Meyers, R. M. et al. Computational correction of copy number effect improves specificity of CRISPR-Cas9 essentiality screens in cancer cells. Nature Genetics 49, (2017).

