## Supplemental Information for "Epithelial-mesenchymal transition is the main driver of intrinsic metabolism in cancer cell lines"

§ To whom correspondence should be addressed:

### Supplementary Tables

| | | Batch Scoring FC | FC Reproducibility | FC Reproducibility AA | Interbatch Distance | Kolmogorov-Smirnov | Average |
|---|---|---|---|---|---|---|---|
| | Not Normalized | 1 | 1 | 1 | 1 | 1 | 1 |
| Batch Effects | RUVs Batch (Batch & MDAMB321) | 0.73 | 0.77 | 0.87 | 1.52 | 0.85 | 0.95 |
| | RUVsOne (MDAMB321) | 0.64 | 0.76 | 0.67 | 1.35 | 0.74 | 0.83 |
| | Combat (Batch) | 1.07 | 0.97 | 1.23 | 0.18 | 1.13 | 0.91 |
| Signal Drift | Moving Median (Plate) | 0.87 | 0.96 | 1.31 | 0.54 | 0.43 | 0.82 |
| | LOESS (Plate) | 0.84 | 0.98 | 0.95 | 0.50 | 0.38 | 0.73 |
| | Robust LOESS (Plate) | 0.84 | 0.96 | 1.40 | 0.54 | 0.41 | 0.83 |
| | QC-SVR (Plate & MDAMB321) | 0.96 | 1.03 | 1.25 | 0.58 | 0.54 | 0.87 |
| Sample Variance | Quantile | 0.15 | 0.17 | 0.05 | 1.32 | 0.03 | 0.34 |
| | Mean | 0.48 | 0.48 | 0.47 | 1.23 | 0.31 | 0.59 |
| | Median | 0.48 | 0.51 | 0.48 | 1.24 | 0.32 | 0.60 |
| | Std | 0.50 | 0.49 | 0.42 | 0.96 | 0.27 | 0.53 |
| | MAD | 0.50 | 0.49 | 0.41 | 0.97 | 0.27 | 0.53 |
| | TIC | 0.48 | 0.48 | 0.46 | 1.17 | 0.30 | 0.58 |
| | PQN | 0.48 | 0.49 | 0.48 | 1.33 | 0.30 | 0.62 |
| | Confluency | 1.13 | 1.00 | 2.63 | 0.68 | 0.94 | 1.28 |
| Combo | MovMedian (Plate) Mean | 0.45 | 0.50 | 0.62 | 0.78 | 0.15 | 0.50 |
| | MovMedian (Batch) Mean | 0.45 | 0.49 | 0.54 | 0.72 | 0.16 | 0.47 |
| | MovMedian (Plate) Combat (Batch) | 0.94 | 0.94 | 1.50 | 0.15 | 0.58 | 0.82 |
| | MovMedian (Plate) Combat (Plate) | 0.94 | 0.91 | 1.31 | 0.24 | 0.54 | 0.79 |
| | Combat (Plate) Mean | 0.46 | 0.46 | 0.79 | 0.32 | 0.19 | 0.45 |
| | Combat (Batch) Mean | 0.51 | 0.47 | 0.62 | 0.33 | 0.22 | 0.43 |
| | MovMedian (Plate) Combat (Batch) Mean | 0.47 | 0.48 | 0.85 | 0.23 | 0.16 | 0.44 |
| | Mean MoMedian (Plate) Combat | 0.48 | 0.47 | 0.76 | 0.22 | 0.19 | 0.42 |
| | Combat (Batch) Quantile | 0.40 | 0.38 | 0.22 | 0.33 | 0.10 | 0.29 |

**Supplementary Table 1**. List of normalization methods and their computed quality metrics score. In parenthesis are the parameter used for normalization, with MDAMB321 one of the control cell line. See details in method.

| Type | Trait | # of Traits | # Significant | Source |
|---|---|---|---|---|
| Experiment | Confluence/Suspension | 2 | 1 | This Study |
| | Batch | 7 | 1 | This Study |
| Metadata | Tissue Type | 11 | 1 | This Study |
| | Pathology | 4 | 0 | CCLE |
| | Histology | 23 | 0 | CCLE |
| | Histology Subtype | 69 | 0 | CCLE |
| | Cancer Type | 38 | 0 | CCLE |
| | Ethnicity | 3 | 0 | CCLE |
| | Mutation rate | 1 | 0 | CCLE |
| Omics | Mutation | 705 | 1 | Li et al. |
| | Copy Number | 61 | 2 | Li et al. |
| | Methylation | 2114 | 252 | Li et al. |
| | Transcript | 56 318 | 453 | CCLE |
| | Protein | 214 | 27 | CCLE |
| Aggregate | Transcription Factor | 743 | 115 | Ortmayr et al. |
| | Signaling Pathway | 14 | 2 | Schubert et al. |
| | Epithelial to Mesenchymal Transition (EMT) | 1 | 1 | Rajapakse et al. |
| TOTAL | | 60 328 | 856 | |

**Supplementary Table 2**. The drivers of metabolic heterogeneity. Associations of a trait to the metabolic phenotype is tested on all main subtrees of the hierarchical clustering tree of the cell lines (obtained in Fig.2 B). A hypergeometric test is used to evaluate association of a qualitative trait to the metabolic phenotype and a student t-test is used evaluate association of a quantitative trait to the metabolic phenotype B. List of all traits integrated to the metabolic phenotype, where traits were consider significant at 10 % FDR (cell lines n = 180).

| Typing | Cell Line | Tissue | Doubling Time | |
|---|---|---|---|---|
| Type 2 | HCT15 | Colon | 20.6 | Adherent |
| Type 2 | NCIH460 | Non-Small Cell Lung | 17.8 | Adherent |
| Type 2 | T47D | Breast | 45.5 | Adherent |
| Type 2 | OVCAR3 | Ovarian | 34.7 | Adherent |
| Type 2 | SW620 | Colon | 20.4 | Adherent |
| Type 1 | SKMEL5 | Melanoma | 25.2 | Adherent |
| Type 1 | OVCAR5 | Ovarian | 48.8 | Adherent |
| Type 1 | HS578T | Breast | 53.8 | Adherent |
| Type 1 | SF539 | CNS | 35.4 | Adherent |

**Supplementary Table 3**. Cell lines selected for validation. Abbreviation: CNS Central Nervous System

Supplementary Figures

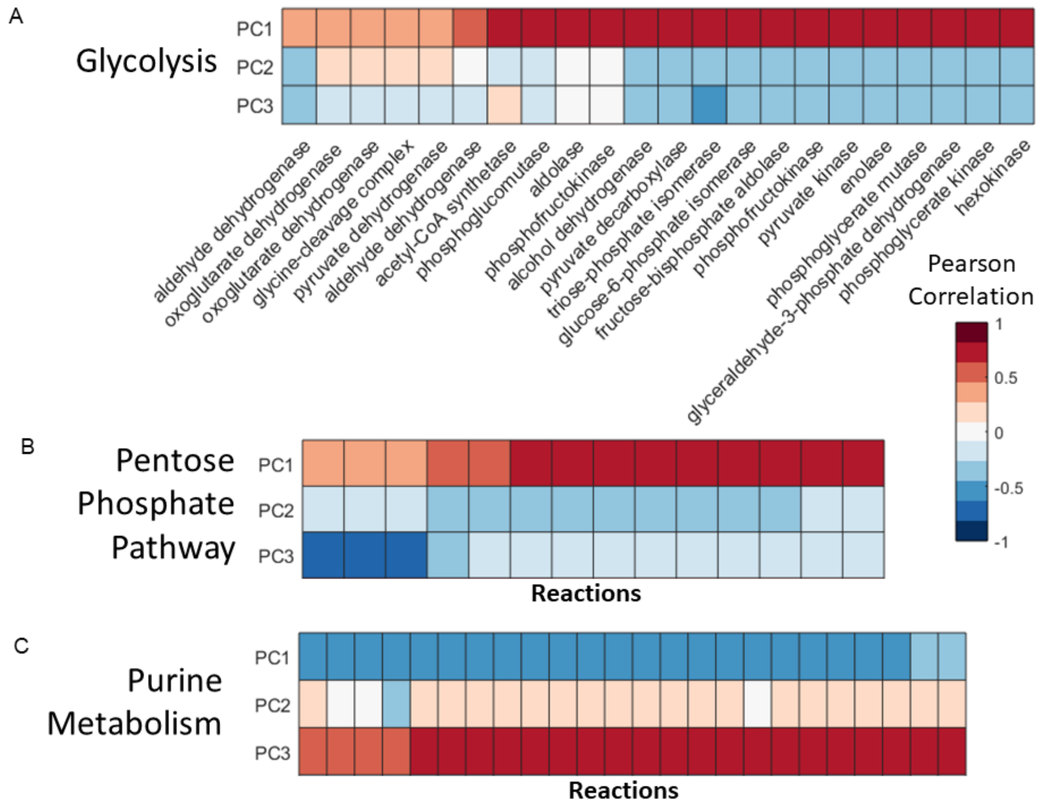


**Figure Sup. 1.** Link between fluxes and metabolites levels. Correlation between fluxes of a pathway and metabolites levels reduces into principal components. For the fluxes and metabolites of A. glycolysis, B. pentose phosphate pathway and C. purine metabolism. Data taken from Hackett et al.[22].

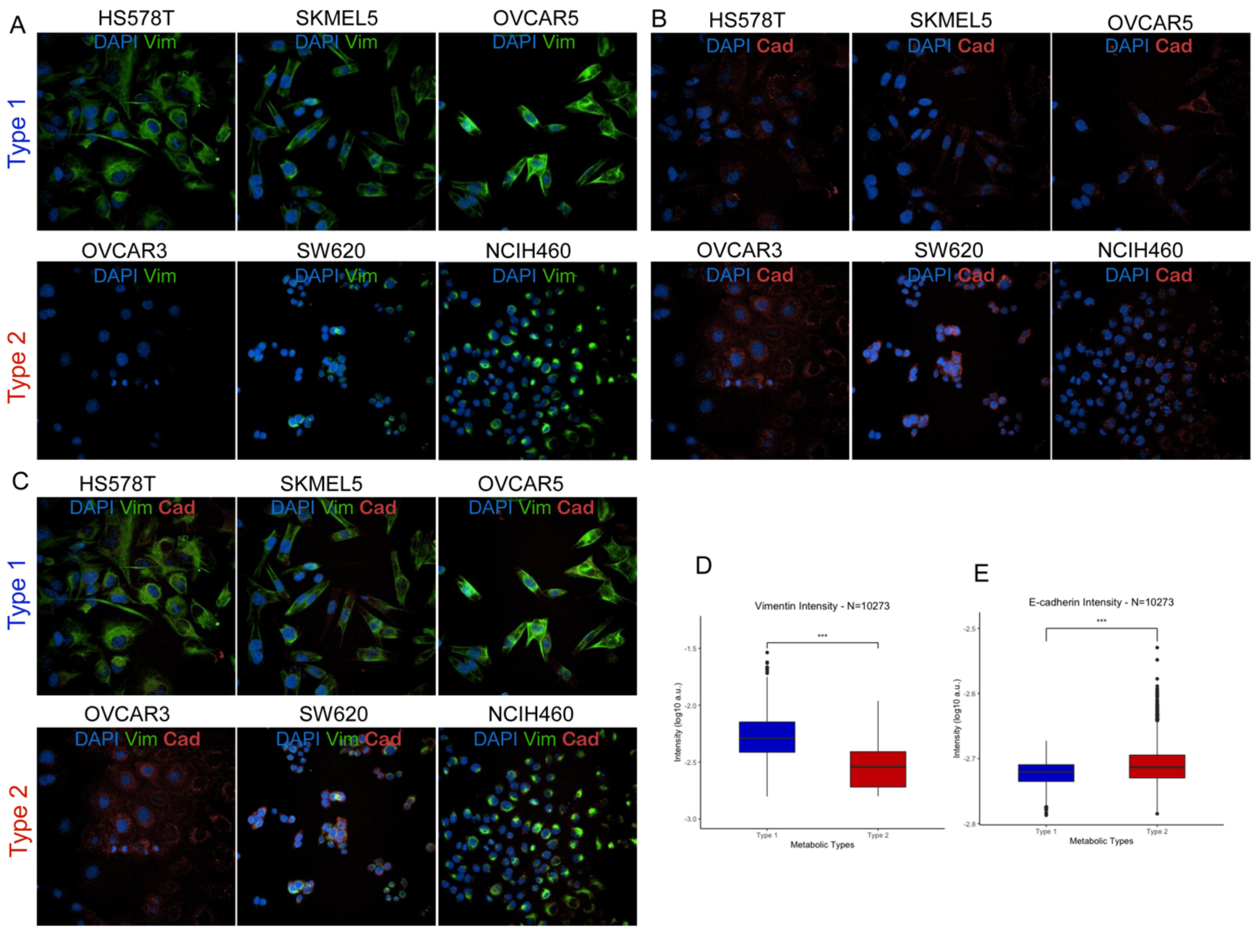


**Supplementary Figure 2.** The metabolic types associated to epithelial and mesenchymal states. Staining examples of the 2 types stained for the nucleus (blue), A. for vimentin (green) and B. e-cadherin (red), C. both, two markers of EMT. Quantification of the expression of D. vimentin and E. e-cadherin in all segmented cell lines (number of cells n = 10 273, two-sided t test) $***p \leq 0.001$; Abbreviation: a.u. arbitrary units

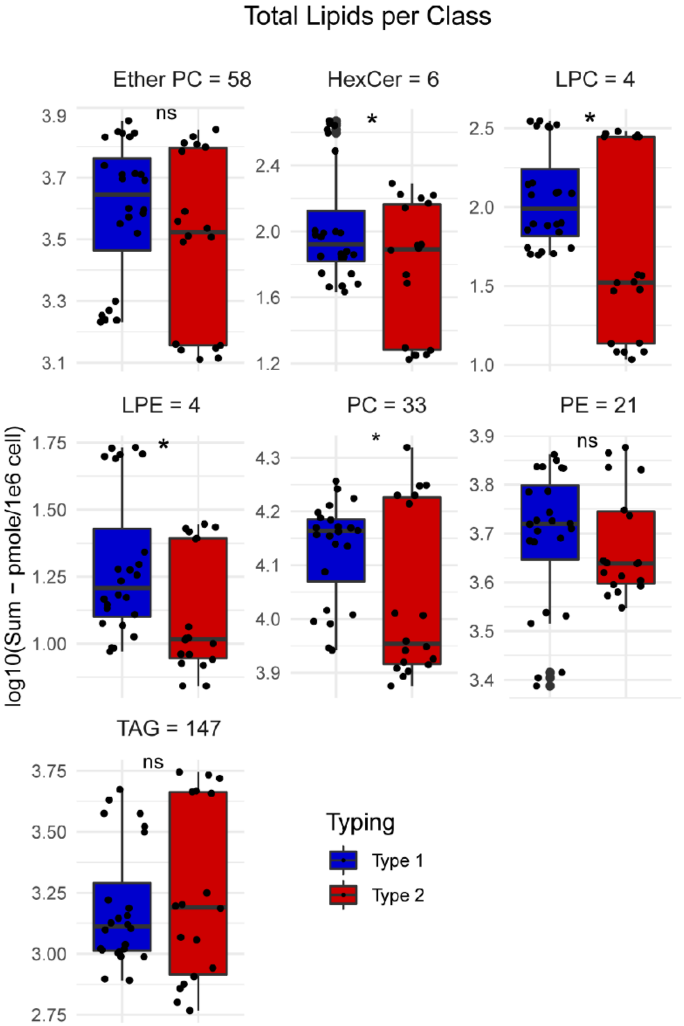


**Supplementary Figure 3.** Differences in lipid content in each class, with number of lipids per class in title (cell lines n=7, Student t-test). ns: p > 0.05, *: p <= 0.05, **: p <= 0.01, ***: p <= 0.001, ****: p <= 0.0001

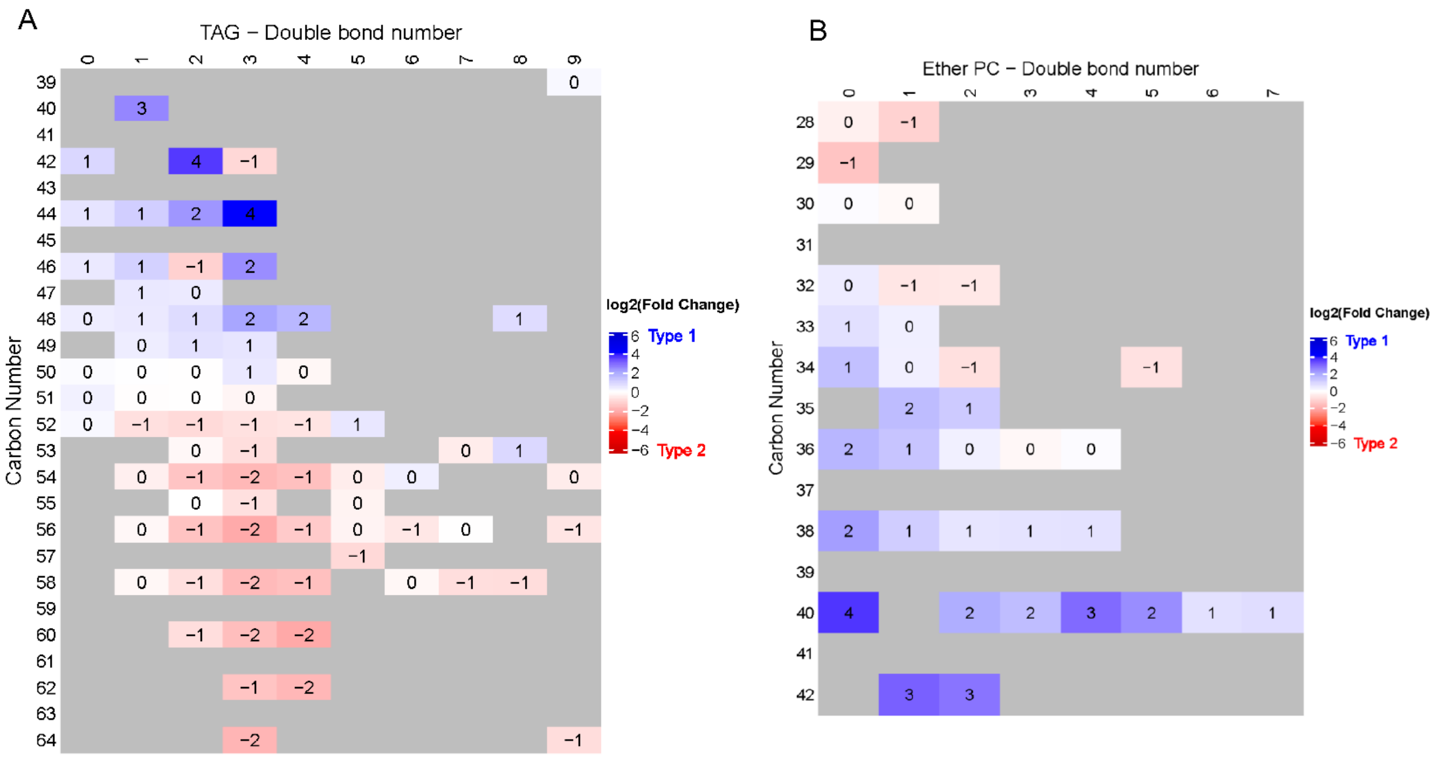


**Supplementary Figure 4.** Differential analysis of all A. TAG and B. Ether PC displayed per double bond number and carbon number.

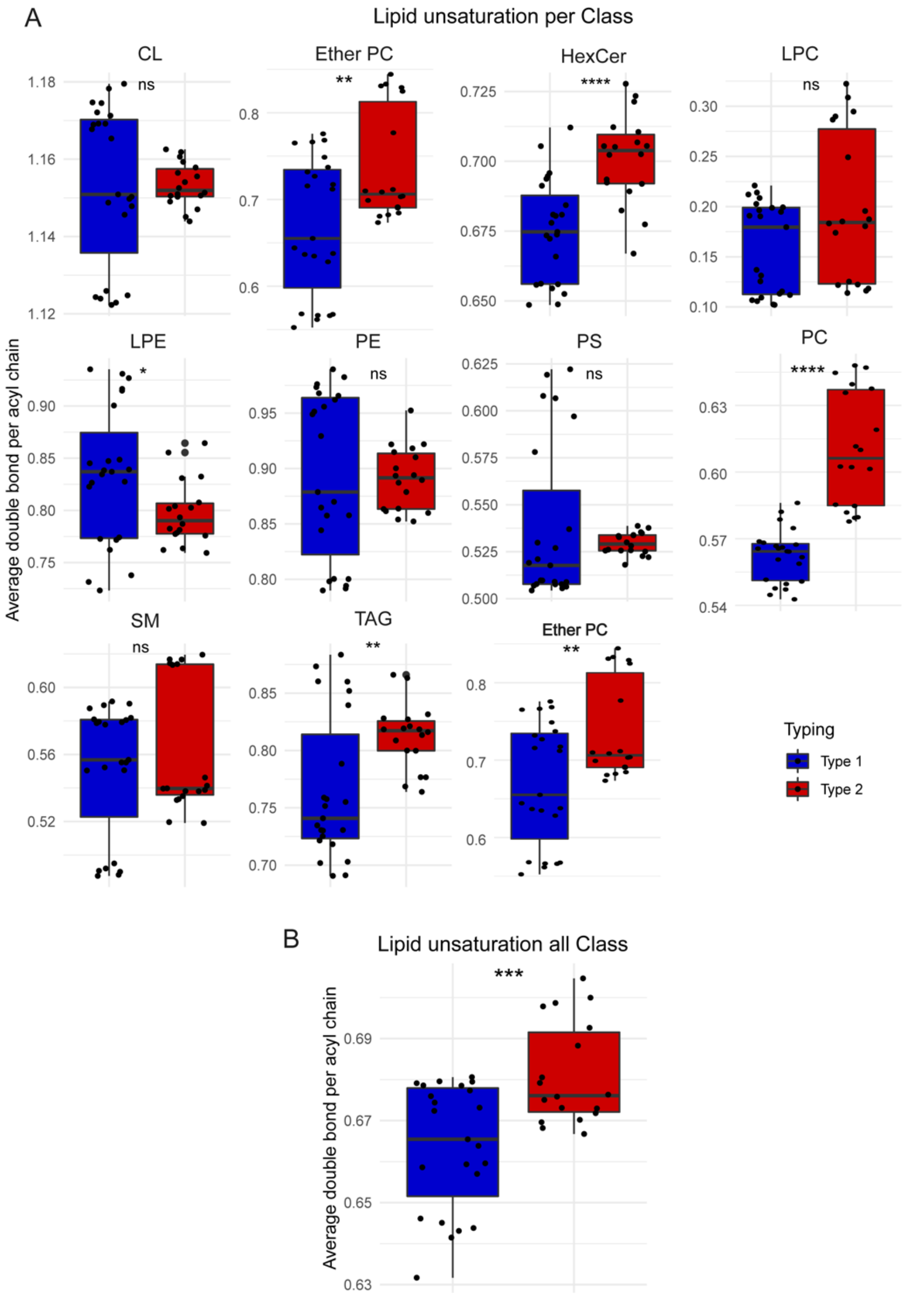


**Supplementary Figure 5**. Difference in unsaturation (concentration-weighted average double per acyl chain) across types (cell lines n=7, Student t-test) for A. lipid class and B. all lipids.

ns: p > 0.05, *: p <= 0.05, **: p <= 0.01, ***: p <= 0.001, ****: p <= 0.0001

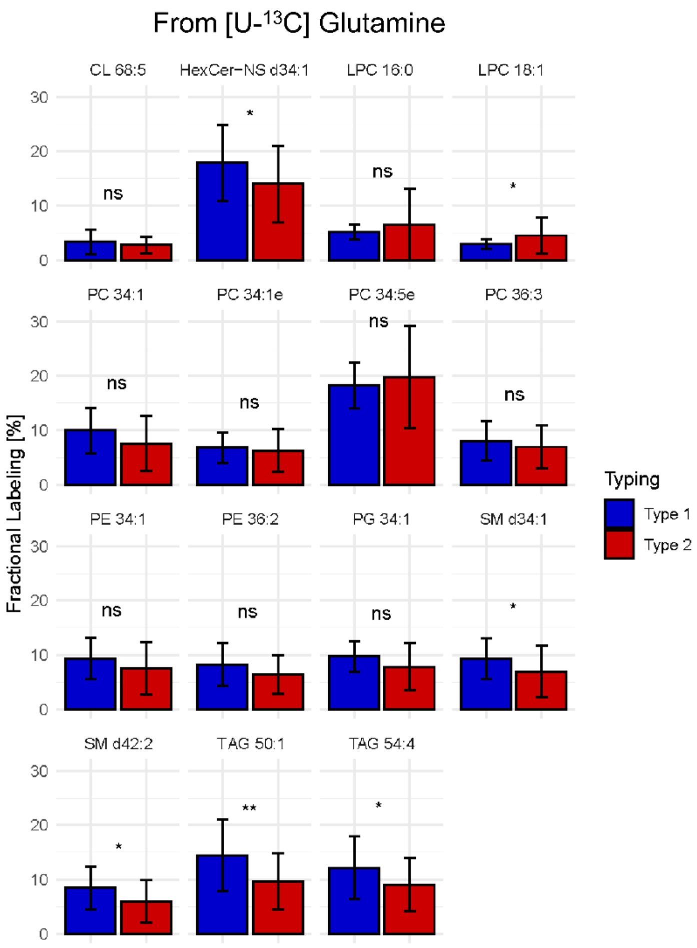


**Supplementary Figure 6.** Differential de novo lipid biosynthesis from [U-$^{13}$C] glutamine. Bar plot (mean +/standard deviation) of fractional labeling from [U-$^{13}$C] glutamine of most abundant lipids per lipid class (cell lines n=9, Student t-test). ns: p > 0.05, *: p <= 0.05, **: p <= 0.01, ***: p <= 0.001, ****: p <= 0.0001

A

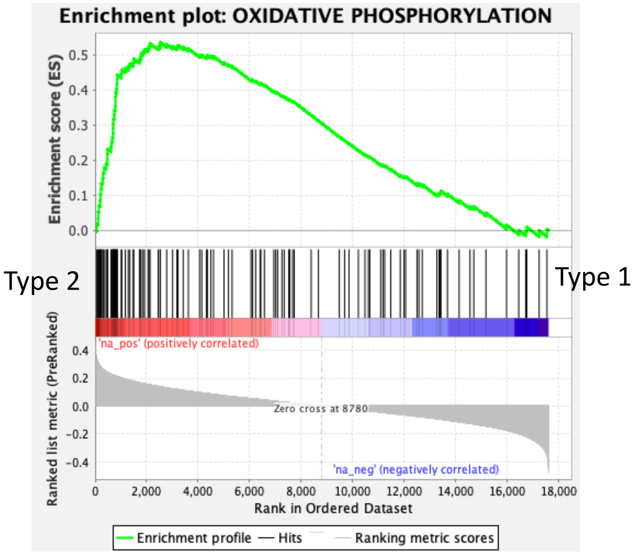


B

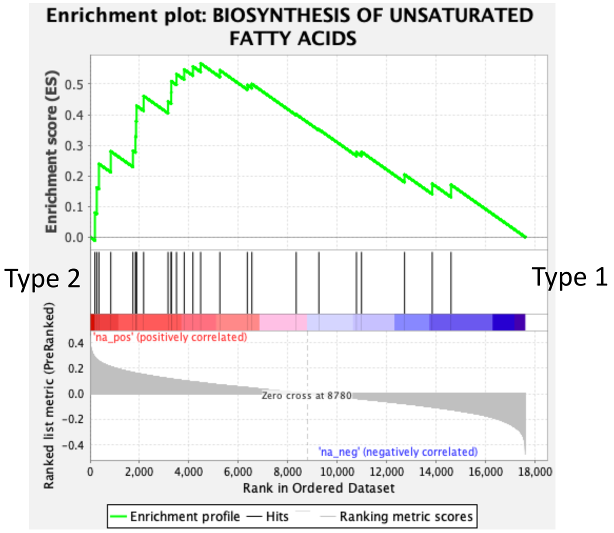


**Supplementary Figure 7.** Pathway dependency of metabolic types. A. Enrichment of oxidative phosphorylation and B. biosynthesis of unsaturated fatty acids. Gene set enrichment analysis result and figures were generated using leading edge analysis described in GSEA (https://www.gsea-msigdb.org/gsea/index.jsp).
